# THRONCAT: Efficient metabolic labeling of newly synthesized proteins using a bioorthogonal threonine analog

**DOI:** 10.1101/2022.03.29.486210

**Authors:** Bob J. Ignacio, Jelmer Dijkstra, Natalia Mora Garcia, Erik F.J. Slot, Margot J. van Weijsten, Erik Storkebaum, Michiel Vermeulen, Kimberly M. Bonger

## Abstract

Profiling the nascent cellular proteome and capturing early proteomic changes in response to external stimuli provides valuable insight into cellular physiology. Existing metabolic protein labeling approaches based on bioorthogonal methionine-or puromycin analogs allow for the selective visualization and enrichment of the newly synthesized proteins. However, their applications are limited as they require methionine-free conditions, auxotrophic cells and/or are toxic to cells. Here, we introduce THRONCAT, a novel threonine-derived non-canonical amino acid tagging method based on bioorthogonal threonine analog β-ethynylserine (βES) that enables efficient and non-toxic labeling of the nascent proteome in complete growth media within minutes. We used THRONCAT for the visualization and enrichment of nascent proteins in bacteria, mammalian cells and *Drosophila melanogaster*. We profiled immediate proteome dynamics of Ramos B-cells in response to receptor activation, demonstrating the ease-of-use of the method and its potential to address diverse biological questions. In addition, using a *Drosophila* model of Charcot-Marie-Tooth peripheral neuropathy, we show that THRONCAT enables visualization and quantification of relative protein synthesis rates *in vivo*.

## INTRODUCTION

Cells rapidly respond to environmental changes by synthesizing new proteins to ensure proper cell functioning. Characterizing this nascent proteome is important to understand cell functioning, but remains challenging as it requires an approach to distinguish between newly synthesized proteins (NSPs) and the pre-existing proteome. Metabolic protein labeling uses exogenous amino acid-or puromycin analogs, which are incorporated into nascent proteins by the endogenous biosynthetic machinery.^1^ These metabolic probes contain bioorthogonal reactive groups for conjugation to fluorescent dyes or affinity tags, enabling selective visualization and enrichment of NSPs (Fig. 1).^2^

**Fig. 1:**
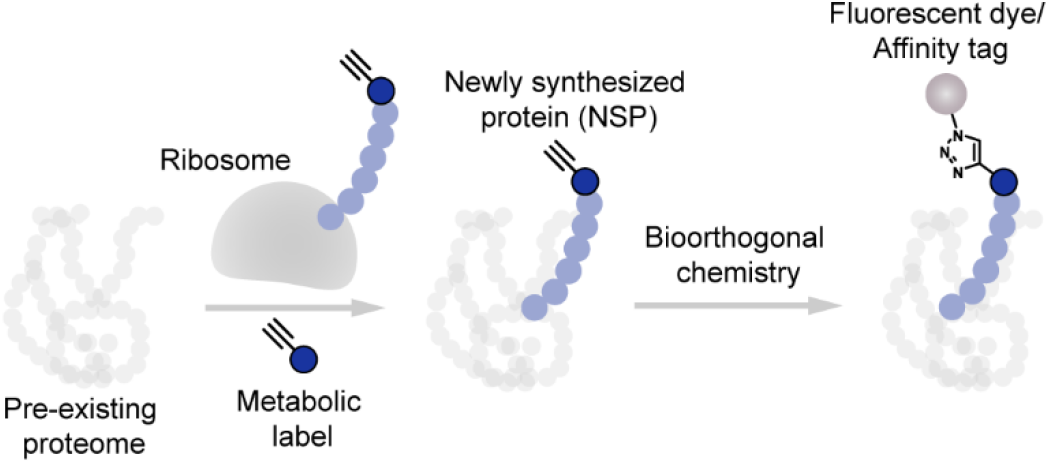
Scheme of metabolic protein labeling. Bioorthogonal analogs of amino acids or puromycin are incorporated biosynthetically into growing polypeptide chains on the ribosome. The metabolic labels are exclusively incorporated into newly synthesized proteins (NSPs), allowing for their selective visualization or enrichment through bioorthogonal ligation to fluorescent dyes or affinity tags, respectively.

The most commonly used metabolic labeling reporters are bioorthogonal methionine analogs azidohomoalanine (AHA) and homopropargylglycine (HPG; Fig. 2a),^3–5^ and puromycin analog *O*-propargyl-puromycin (OPP).^6^ OPP labels NSPs within minutes and is easy to use, as labeling can be performed in complete growth media. However, OPP is toxic to cells and yields truncated OPP-polypeptide adducts that are unstable and proteolytically degraded within 1 hour.^6^ In contrast, bioorthogonal non-canonical amino acid tagging (BONCAT) with AHA and HPG is less toxic and provides stable labeling in NSPs. However, as AHA and HPG are poorly incorporated in proteins in the presence of methionine, BONCAT requires methionine starvation, precluding its use to study protein synthesis dynamics in their native context.^7^

**Fig. 2:**
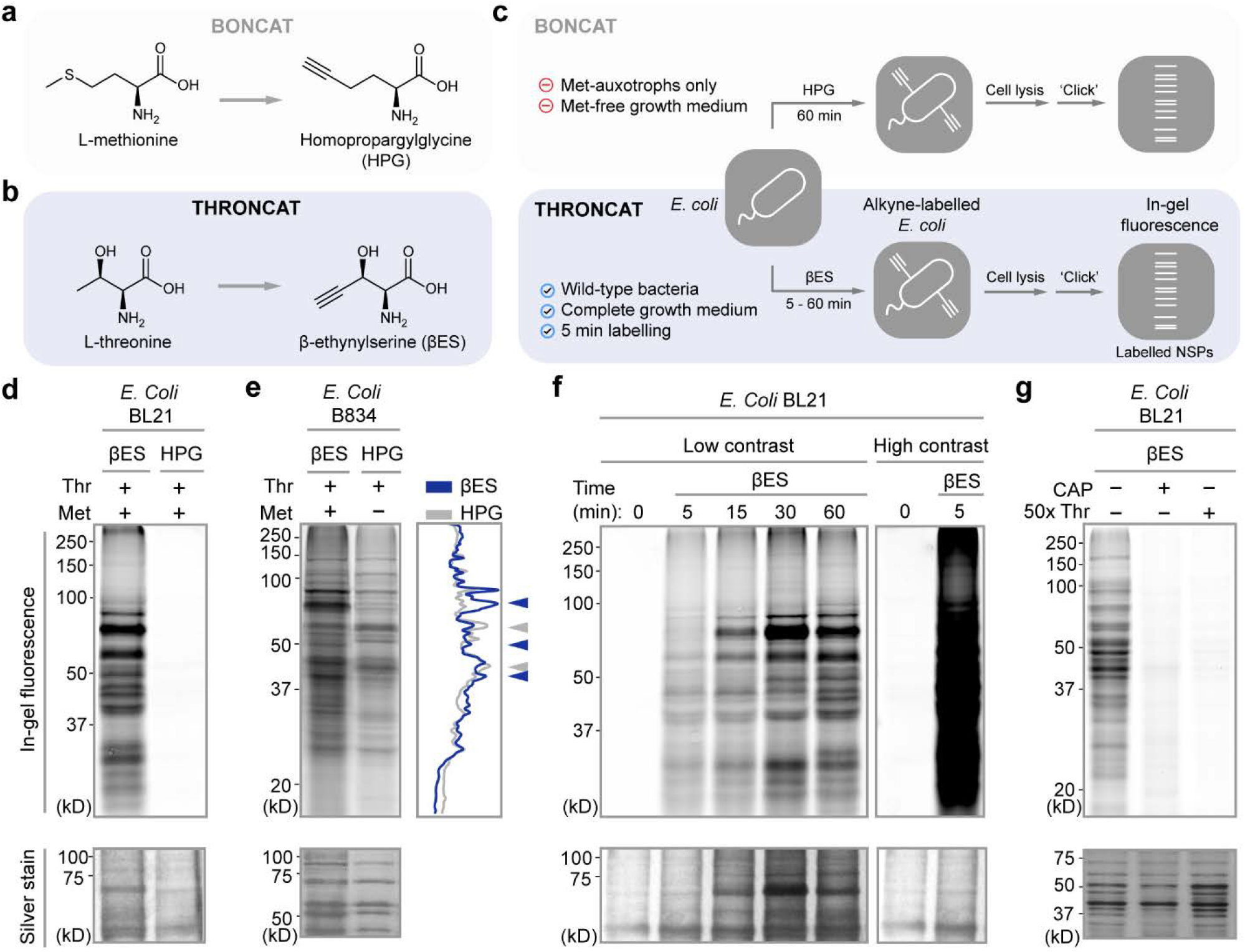
Characterization of βES labeling in bacteria. **a**, Structures of L-methionine and its analog homopropargylglycine (HPG) used in BONCAT. **b**, Structures of L-threonine and its analog β-ethynylserine (βES) used in THRONCAT. **c**, Scheme of the BONCAT and THRONCAT workflows for NSP labeling in *E. coli* and their associated drawbacks and benefits. **d-g**, In-gel visualization of βES or HPG incorporated into *E. coli*. Incorporated βES or HPG was conjugated to Cy5-azide using a copper-catalyzed azide-alkyne cycloaddition reaction and visualized through in-gel fluorescence scanning. Silver stain panels show total protein. **d**, Comparison of βES and HPG incorporation into *E. coli* BL21 lysate after 1 h incubation with 4 mM βES or 4 mM HPG in LB medium. **e**, Detection of βES and HPG in methionine auxotrophic *E. coli* B834 lysate after 1 h incubation with 4 mM of βES or 4 mM HPG in Selenomet medium. Line scan of gel lanes shows differences in labeling intensities between corresponding bands in βES (blue line) and HPG (gray line) treated samples. Arrows indicate selected bands showing strong relative labeling in βES (blue arrows) and HPG (grey arrows) treated samples. Met, Methionine. **f**, βES incorporation in *E. coli* BL21 over time. *E. coli* was incubated with 4 mM βES in LB medium for the indicated durations. ‘High contrast’ panel is a duplicate of the 0 min and 5 min time points in the ‘low contrast’ panel, but with enhanced contrast to highlight the large difference in labeling intensities. **g**, βES detection in *E. coli* BL21 after 1 h incubation with 1 mM βES in LB medium. No βES incorporation is detected upon co-incubation with chloramphenicol or an excess of threonine. CAP, chloramphenicol; Thr, L-threonine; kD, Kilodalton.

These challenges highlight the need for a metabolic protein labeling method that allows fast and efficient incorporation of amino acid analogs in nascent proteins and can be used under native conditions. Threonine analogs 4-fluorothreonine (4-FT)^8^ and β-hydroxynorvaline (β-HNV)^9,10^ are excellent substrates for the threonine tRNA synthetase (ThrRS) and are efficiently incorporated into the nascent proteome of bacteria in complete growth media. We envisioned that the bioorthogonal threonine analog β-ethynylserine (βES; Fig. 2b),^11–13^ may also be efficiently incorporated into nascent proteins, facilitating their labeling in cells grown in complete medium.

Here, we introduce THRONCAT, a novel metabolic labeling method based on <u>thr</u>e<u>o</u>nine-derived <u>n</u>on-<u>c</u>anonical <u>a</u>mino acid tagging. We show that threonine analog βES is efficiently incorporated into NSPs, is non-toxic and allows labeling of the nascent cellular proteome in complete growth medium within minutes. We demonstrate that THRONCAT has high labeling efficiency and compare our method to BONCAT for the visualization of NSPs in bacteria, mammalian cells and *Drosophila melanogaster*. Furthermore, we used THRONCAT to profile proteomic changes over time using βES pulse-labeling following B-cell receptor (BCR) stimulation in Ramos B-cells, demonstrating the accessibility and ease-of-use of the method. In addition, by combining THRONCAT with genetic expression of a fluorescent cell marker in motor neurons, we quantified *in vivo* cell type specific changes in protein synthesis rate in a *Drosophila* model of Charcot-Marie-Tooth peripheral neuropathy.

## RESULTS

### βES is efficiently incorporated into the nascent prototrophic *E. coli* proteome

To explore the use of threonine analogs in metabolic labeling experiments, we first devised a stereoselective synthesis route towards βES (Supplementary Fig. 1). Starting from commercially available materials and using an asymmetric aminohydroxylation as key step in our synthetic route, we obtained βES in four steps in a 22% overall yield (>98% *de*, 86% *ee*).

Next, we aimed to explore whether βES is incorporated into the nascent proteome of bacteria. As prototrophic bacteria can synthesize their own pool of methionine, tagging NSPs using methionine derivatives is generally challenged by the need of methionine-auxotrophic bacteria and the use of methionine-free growth medium. Indeed, we observed no labeling using methionine derivative HPG in prototrophic *E. coli* BL21 cells that were grown in complete growth medium (Fig. 2c, d), while we confirmed strong incorporation of HPG in auxotrophic *E. coli* B834 cells in methionine-free growth medium (Fig. 2e). In contrast to HPG, supplementing prototrophic *E. coli* BL21 cells growing in complete medium with βES resulted in strong labeling of the nascent proteome (Fig. 2c,d). The labeling ensued throughout the proteome, in a time-dependent manner and without significantly affecting bacterial growth (Fig. 2d, f; Supplementary Fig. 2). Incorporation of βES was abrogated by the addition of protein synthesis inhibitor chloramphenicol (CAP) or a 50-fold excess of threonine, suggesting that βES is incorporated exclusively into NSPs at positions encoding for threonine (Fig. 2g). Interestingly, looking at the relative labeling intensities of individual protein bands in the SDS-PAGE gels, we observed a clear difference between βES-and HPG-labeled *E. coli* lysate, possibly because of varying numbers of threonine- and methionine residues in individual proteins (Fig. 2e).

### Using βES for fast and non-toxic visualization of mammalian NSPs

Encouraged by the observation that βES is vastly incorporated in NSP of prototrophic *E. coli* in complete medium, we next explored the efficiency of βES labeling of nascent proteins in mammalian cells. We treated HeLa cells for 1 h with βES in complete growth medium and conjugated incorporated βES to a Cy5-azide fluorophore. Fluorescence analysis of HeLa cells by flow cytometry and in-gel fluorescence revealed efficient and concentration-dependent incorporation of βES (Fig. 3a,b; Supplementary Fig. 3). Co-incubation with protein synthesis inhibitor cycloheximide or excess threonine abolished all fluorescence, confirming incorporation of βES at the position of threonine (Supplementary Fig. 4). The fluorescence signal was discernible from background using the lowest (4 μM βES) concentration tested and increased dose-dependently, giving a ∼200-fold increase in signal over background at the highest concentration tested (4 mM βES; Fig. 3a). In contrast, 1 h incubation of HeLa cells with 4 mM HPG in complete medium yielded only minimal HPG incorporation in NSPs (Fig. 3a, Fig. 3b). Importantly, we could obtain an additional 1000-fold increase in detection sensitivity using βES when cells were grown in medium that was depleted from threonine (*e*.*g*. 200-fold signal over background at 4 μM βES, Supplementary Fig. 5).

**Fig. 3:**
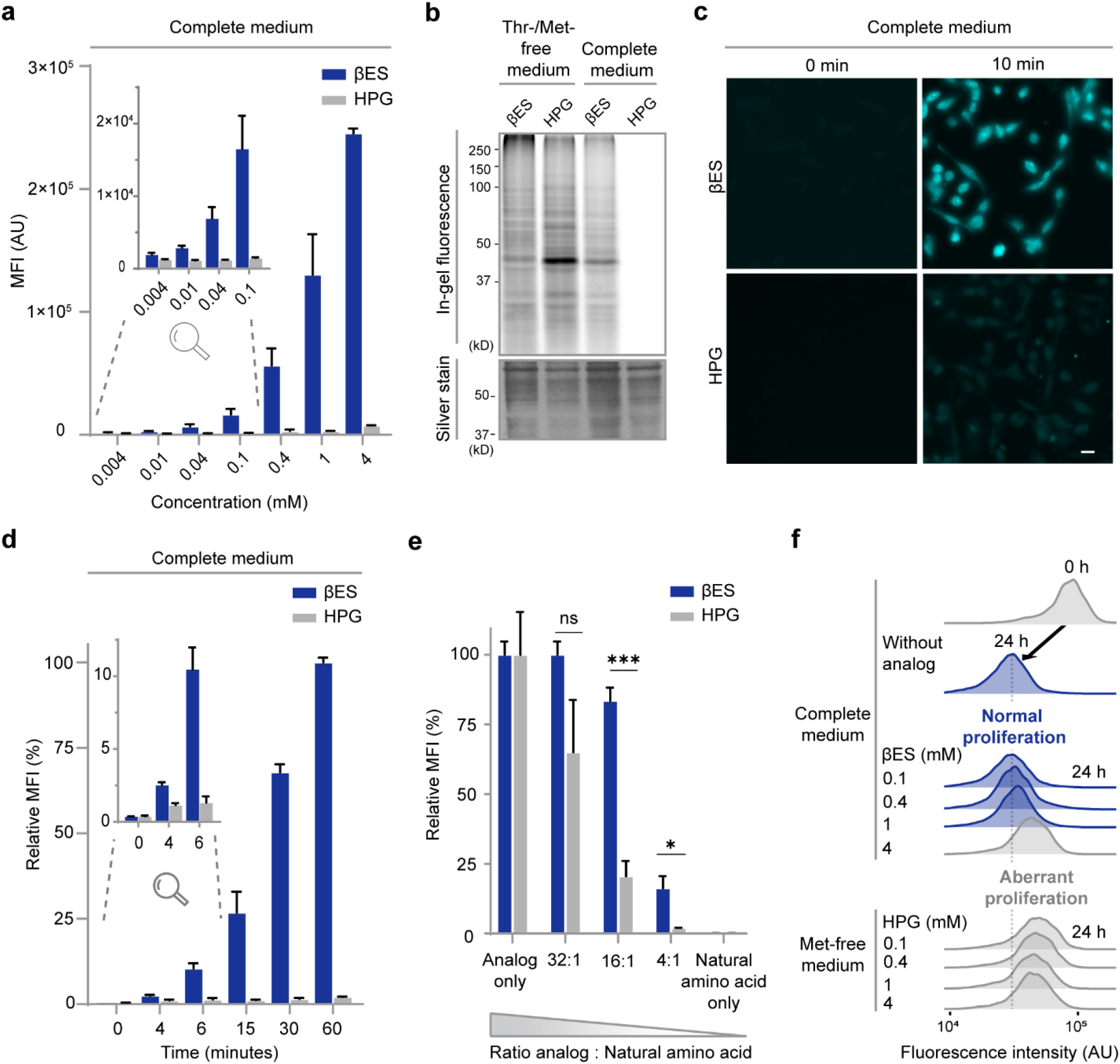
Visualization of NSPs in mammalian cells using THRONCAT. **a-e**, Visualization and quantification of incorporated analogs was enabled by conjugation to Cy5-azide. **a**, Flow cytometry quantification of βES or HPG incorporation into HeLa cells. HeLa cells were incubated for 1 h with the indicated concentration of analogs in complete medium. Top-left inset shows zoom of 0.004 – 0.1 mM concentrations. MFI, mean fluorescence intensity; AU, arbitrary units. Error bars represent s.d. Sample size is *n* = 3. **b**, In-gel detection of βES or HPG in HeLa proteome after 1 h incubation with analog. HeLa cells were labeled with 4 mM βES or 4 mM HPG in threonine-free or methionine-free medium, respectively, or in complete medium. Silver stain panel shows total protein. **c**, Representative fluorescent images of HeLa cells treated for 10 min with 4 mM βES or 4 mM HPG in complete medium. Untreated HeLa cells (0 min) were also subjected to conjugation with Cy5-azide. Scale bar, 20 μm. **d**, Flow cytometry time-course analysis of βES and HPG incorporation into HeLa proteomes. HeLa cells were incubated with 4 mM analog in complete medium for the indicated duration. Top-left inset shows zoom of time points 0 – 6 min. Signals are normalized to that of βES at 60 min, which is set to 100%. Error bars represent s.d. Sample size is *n* = 3. **e**, Flow cytometry quantification of the inhibitory effect of the natural amino acids, threonine or methionine, on the incorporation of their analog, βES or HPG, respectively. HeLa cells were incubated with 50 μM analog and increasing ratios of threonine or methionine, respectively. For both analogs, signals are normalized to their respective signals for ‘analog only’, which are set to 100%. Ns, not significant. ***P = 0.0001 and *P = 0.0305 (95% confidence interval); determined by an unpaired two-tailed *t* test. Error bars represent s.d. Sample size is *n* = 3. **f**, Flow cytometry proliferation assay of HeLa cells. HeLa cells were labeled with Celltrace Violet and incubated without analog (control) or with indicated concentrations of analog for 24 h. Black arrow indicates decrease in Celltrace fluorescence in control cells. Dotted line indicates final fluorescent signal obtained from control cells. Conditions giving normal proliferation are indicated in blue and conditions giving aberrant proliferation are indicated in grey.

Fluorescence microscopy and flow cytometry revealed that labeling with 4 mM βES in complete medium gave a strong fluorescent labeling of the nascent HeLa proteome after Cy5-azide conjugation within minutes (Fig 3c, d). The NSPs were distributed throughout the whole cell and strongest fluorescence was observed in the nucleoli (Fig. 3c), which is consistent with previous results obtained with HPG and OPP labeling.^6^

The results above suggest that βES incorporation is less sensitive to competition of threonine than HPG for methionine. Indeed, while competition of threonine decreased βES incorporation in HeLa cells, we observed a much stronger inhibitory effect for methionine on HPG incorporation (Fig. 3e).

Next, we assessed the incorporation of βES over longer time periods. We incubated HeLa cells up to 24 h with βES in complete medium and observed a steady increase of βES incorporation as quantified by flow cytometry (Supplementary Fig. 6). Importantly, we did not observe a reduction in cell viability at 24 h in the presence of 0.4 – 4 mM βES as measured by propidium iodide exclusion (Supplementary Fig. 7). In addition, to reveal any adverse effects on cell proliferation, we observed no effect of protein labeling on HeLa cells exposed to up to 1 mM βES for 24 h. Incubation of cells with higher βES concentration, however, resulted in slightly decreased proliferation rate compared to control cells, which we also observed in HeLa cells treated with the lowest concentration HPG tested (e.g. 0.1 mM) for 24 h in methionine-free medium (Fig. 3f).

### THRONCAT enables identification and quantification of the dynamic proteome

Next, we were interested to explore THRONCAT in the enrichment and detection of NSPs of HeLa cells and compared the nature of the identified proteins with that observed when using BONCAT in a mass-spectrometry-based proteomics setup. To this end, we incubated HeLa cells in complete medium for 5 h with 4 mM βES in 3 biological replicates, enriched NSPs and subjected digested peptides to LC-MS/MS analysis (Fig. 4a). By selecting for proteins that were identified in all 3 biological replicates, we confidently identified 3073 unique HeLa NSPs (Fig. 4b, Supplementary Data 1). Comparison of the individual replicates shows that ∼ 83% of NSPs is found in all 3 replicates and ∼ 90% by at least 2 replicates, indicating good experimental reproducibility (Supplementary Fig. 8). Using an original Python script (Supplementary Data 2), we determined that the average threonine content of identified proteins is 5.20%, which is identical to the threonine content of the human proteome showing that THRONCAT does not enrich for threonine-rich proteins. Labeling with 1 mM βES identified 2751 NSPs, only a ∼ 10% reduction, highlighting the efficiency of βES incorporation into the nascent proteome (Supplementary Fig. 9). Using BONCAT – 4 mM HPG in methionine-free conditions – we identified a similar number of HeLa NSPs as with 4 mM βES in complete medium (Fig. 4b). Overlap between NSPs identified by THRONCAT and BONCAT was large (81%) and only a fraction of NSPs were identified solely by THRONCAT (9.4%) or BONCAT (9.5%) (Fig. 4c). Interestingly, the THRONCAT-only fraction was enriched in proteins low in methionine content and conversely, the BONCAT-only fraction was enriched with proteins low in threonine content (Supplementary Fig. 10, Supplementary Data 3).

**Fig. 4:**
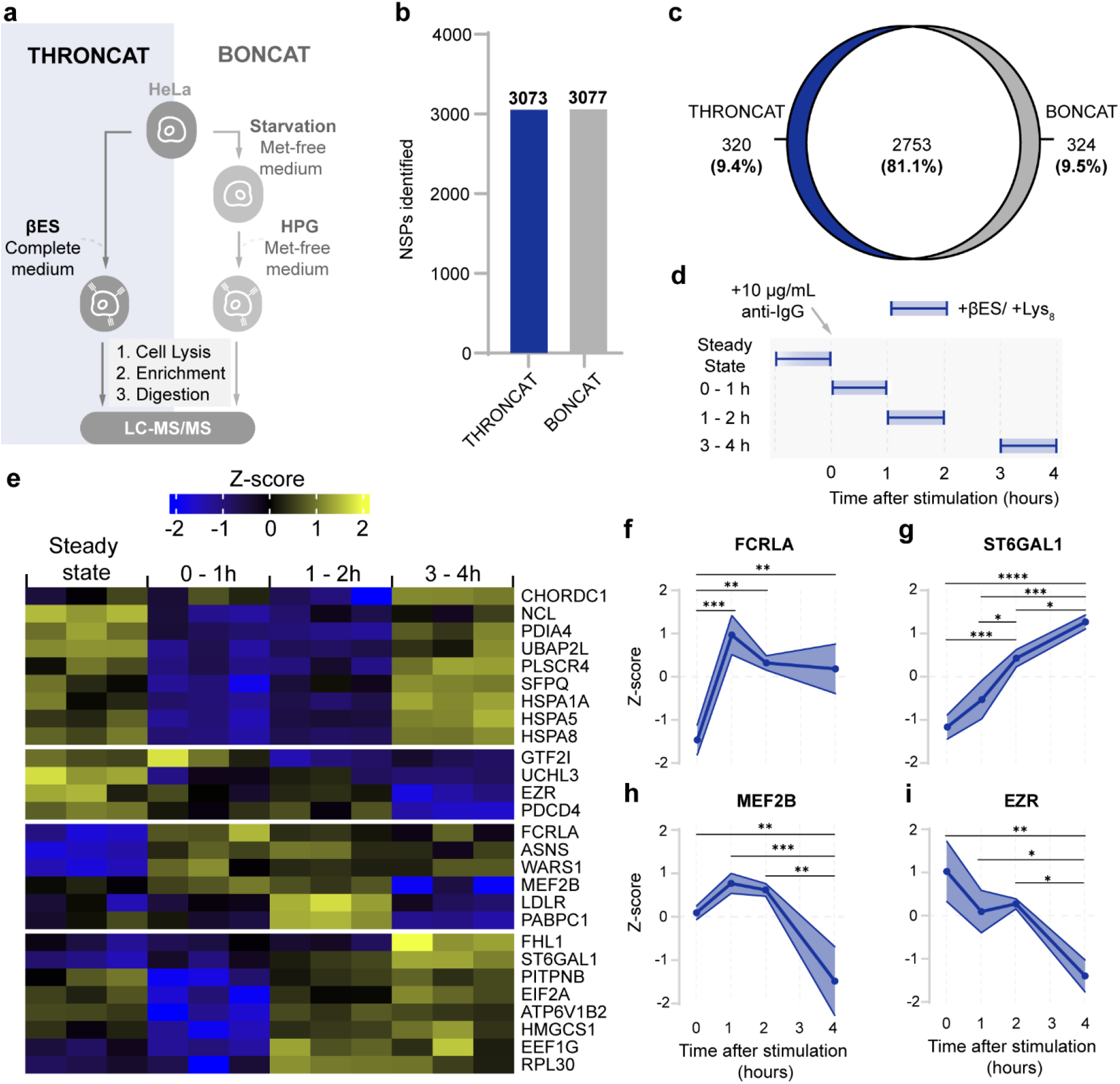
Proteomic analysis of the nascent mammalian proteome with THRONCAT. **a**, Scheme showing THRONCAT and BONCAT workflows for proteomic analysis of NSPs. NSP, newly synthesized protein; LC-MS/MS, Liquid chromatography-tandem mass spectrometry. **b**, Number of HeLa NSPs confidently identified by THRONCAT and BONCAT. HeLa cells were incubated for 5 h with 4 mM βES in complete medium (THRONCAT) or starved in methionine-free medium and then incubated with 4 mM HPG in methionine-free medium for 5 h (BONCAT). Cells were lysed and NSPs were selectively enriched, digested and their peptides subjected to LC-MS/MS analysis. Untreated HeLa cells were used as a control. Experiment was performed with three biological replicates per condition. NSPs were assigned as confident if they appeared in all three replicates and did not appear in all three replicates of the untreated control. **c**, Venn diagram showing overlap between NSPs identified with THRONCAT and BONCAT. **d**, Schematic overview of time course pulse-labeling of stimulated Ramos B cells using THRONCAT. Blue boxes indicate pulse-labeling time windows during which Ramos cells were exposed to 1 mM βES and d_8_-lys. Ramos cell activation was initiated by addition of 10 μg/mL anti-IgG at *t* = 0. Experiment was performed with three biological replicates per time window. Anti-IgG, Anti-immunoglobin G antibody; lys_8_, ^*13*^C_6_,^15^N_2_-*L-*lysine. **e**, Heatmap of differentially expressed proteins after Ramos cell activation. Enriched NSPs were quantified by label-free quantification and their abundances normalized by z-score. Only identified proteins that were present in all replicates of all-time windows and had an adjusted *p* < 0.05 and fold change > 1.5 are included. **f-i**, Expression profiles of FCRLA, ST6GAL1, MEF2B and EZRIN constructed with z-scores from the heatmap shown in **e**. Significance determined by one-way ANOVA, with Bonferroni correction for multiple testing. **f**, FCRLA, Fc receptor-like A. From left to right, *\*\*\*P =* 0.004, *\*\*P* = 0.0033, *\*\*P =* 0.0054. **g**, ST6GAL1, B-galactoside alpha-2,6-disialyltransferase 1. From left to right, *\*\*\*\*P* < 0.0001, *\*\*\*P* = 0.0007, *\*P =* 0.0162, *\*\*\*P =* 0.0003, *\*P =* 0.0324. **h**, MEF2B, Myocyte enhancer binding factor 2B. From left to right, *\*\*P* = 0.0083, *\*\*\*P* = 0.0009, *\*\*P =* 0.0014. **i**, EZR, Ezrin. From left to right *\*\*P* = 0.0027, *\*P* = 0.0258, *\*P* = 0.0131.

Because of the fast kinetics and easy workflow, we envisioned that THRONCAT may enable capturing immediate protein abundance dynamics following cell stimulation by pulse labeling cells at different time intervals. For this, we stimulated Ramos B-cells with anti-IgG and pulse-labelled cells for 1 h at different time points with 1 mM βES as well as d_8_-lysine, which provided a unique internal marker for NSPs. Measuring 3 biological replicates, we pulse-labelled 0–1 h, 1–2 h and 3–4 h after B-cell stimulation and included a pre-stimulation control representing the steady state proteome (Fig. 4d).

Interestingly, using THRONCAT we detected very diverse expression profiles for differentially expressed proteins over the course of B-cell stimulation. We found that of the 579 proteins identified during steady state conditions, 169 (∼ 32%) are not found 0-1 h after anti-IgG stimulation and we detected 114 (∼ 20%) of these proteins again at 1-2 h or 3-4 h after stimulation, suggesting their expression is only temporarily down-regulated upon B-cell activation (Supplementary Data 4; Supplementary Fig. 11, 12). Of the 656 proteins identified, we found 27 differentially expressed proteins (P < 0.05, fold change > 1.5) (Fig. 4e). For instance, overexpression of FCRLA – an intracellular B-cell protein with Fc-receptor binding properties^14^ – reaches peak levels 0-1 h following B-cell activation, (Fig. 4f) while expression of sialyltransferase STGAL6^15^ increases continuously for 4 h (Fig. 4g). Similarly, we observed initial up-regulation of transcription factor MEF2B^16^ followed by down-regulation after 2 h (Fig. 4h), while expression of structural protein ezrin^17^ decreased continuously on the timescale measured (Fig. 4i).

### THRONCAT allows in vivo analysis of protein synthesis in *Drosophila melanogaster*

We next explored whether βES is incorporated into NSPs in *Drosophila melanogaster*, a living and behaving model organism with a rich genetic tool kit. We fed second instar *Drosophila* larvae that selectively express a membrane-tethered green fluorescent protein in motor neurons (OK371-GAL4>UAS-mCD8::GFP) for 48 h on *Drosophila* medium containing 4 mM βES (Fig. 5a). Following labeling, the larval central nervous system (CNS) was dissected and subjected to conjugation chemistry with TAMRA-azide. Although βES is expected to be incorporated in all cells in the body, we decided to focus on the ventral nerve cord (VNC, equivalent to the vertebrate spinal cord) and in particular on motor neurons because of the relevance of protein synthesis (defects) in this cell type for neuromuscular diseases. Confocal microscopy revealed robust THRONCAT labeling in VNC of larvae incubated with 4 mM βES compared to an untreated control (Fig. 5b,c). We observed that βES incorporation was strongly concentration dependent, decreasing ∼ 90% at a 10-fold lower concentration of βES (0.4 mM) (Fig. 5d).

**Fig. 5:**
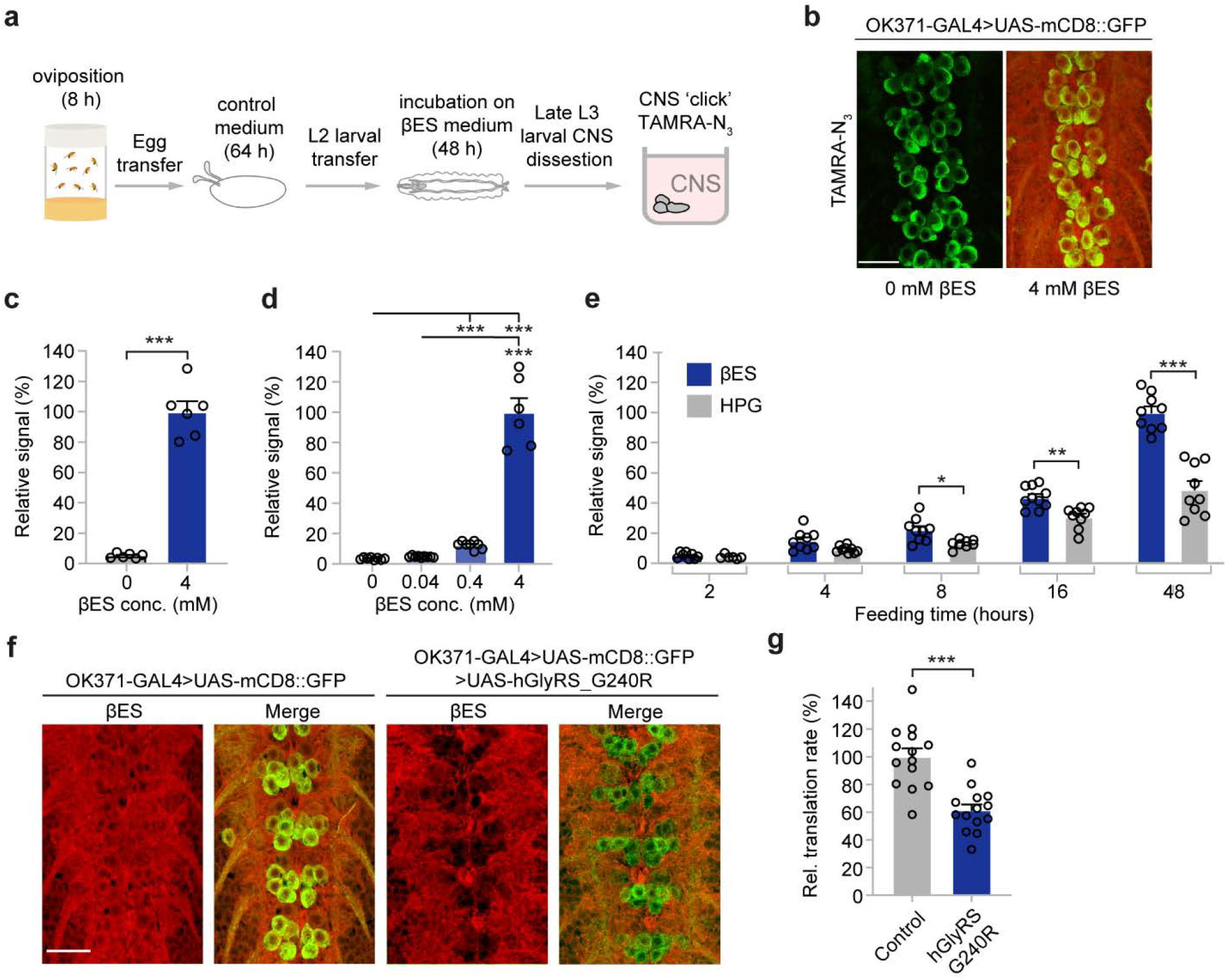
THRONCAT allows *In vivo* analysis of protein synthesis in *Drosophila melanogaster*. **a**, Schematic overview of THRONCAT workflow in Drosophila larvae. CNS, central nervous system; TAMRA-N_3_, 5-carboxytetramethylrhodamine-azide. **b**, Representative images of *in vivo* THRONCAT in *Drosophila* larvae selectively expressing membrane-tethered GFP in motor neurons (OK371-GAL4>UAS-mCD8::GFP). Larvae were exposed to medium containing either 0 mM or 4 mM βES, followed by conjugation to TAMRA-N_3_. Merged images of the green (GFP) and red (TAMRA) channels are shown. Scale bar: 20 μm. **c**, Relative fluorescent signal intensity in larvae exposed to 0 mM or 4 mM βES for 48 h and subjected to TAMRA-N_3_ conjugation. Signals are normalized to that of 4 mM βES, which is set to 100%. *n* = 6-7 larvae per concentration; \*\*\**P* < 0.005 by Mann-Whitney test. Error bars represent SEM. **d**, Relative fluorescent signal intensity in larvae exposed to 0, 0.04, 0.4, or 4 mM βES for 48 h, after conjugation to TAMRA-N_3_. Signals are normalized to that of 4 mM βES, which is set to 100%. *n* = 6-9 larvae per concentration; \*\*\**P* < 0.005 by Kruskall-Wallis test. Error bars represent SEM. **e**, Relative fluorescent signal intensity after conjugation to TAMRA-N_3_ in larvae exposed to 4 mM βES or 4 mM HPG for 2 h, 4 h, 8 h, 16 h, or 48 h. *n* = 6-10 larvae per treatment group and time point; \**P* < 0.05, \*\**P* < 0.01; \*\*\**P* < 0.0005 by unpaired *t*-test (2 h, 16 h, 48 h) or Mann-Whitney test (4 h, 8 h) per time point, with Bonferroni correction for multiple testing. See Table S1 for exact *p*-values. Error bars represent SEM. **f**, Representative images of THRONCAT in Drosophila larvae selectively expressing membrane-tethered GFP in motor neurons (OK371-GAL4>UAS-mCD8::GFP), with or without co-expression of G240R mutant human glycyl-tRNA synthetase (UAS-hGlyRS_G240R). Larvae were exposed to 4 mM βES for 48 h, followed by conjugation to TAMRA-N_3_. Merged images of the green (GFP) and red (TAMRA) channels are shown. Scale bar: 20 μm. **g**, Relative translation rate in motor neurons (OK371-GAL4) as determined by THRONCAT in larvae expressing hGlyRS_G240R versus driver-only control. Signals are normalized to that of control, which is set to 100%. *n* = 14 larvae per genotype; \*\*\**P* < 0.0001 by unpaired *t*-test. Error bars represent SEM.

Next, we investigated the *in vivo* labeling kinetics of βES in comparison to HPG by incubating larvae for different durations with either analog. Signal intensity increased significantly over time for both analogs, but increased faster for βES and reached a higher maximum level (Table S1, Fig. 5e). Notably, 16h exposure to βES resulted in a comparable signal intensity as 48h exposure to HPG, illustrating that THRONCAT can be used to efficiently label NSPs *in vivo* within shorter time frames.

Finally, we determined whether *in vivo* THRONCAT can be used to quantify changes in protein synthesis rates. Using *in vivo* cell-type-specific MetRS^L262G^-ANL FUNCAT,^18^ we previously showed that expression of human glycyl-tRNA synthetase (GlyRS) carrying mutations that cause Charcot-Marie-Tooth (CMT) peripheral neuropathy reduce global protein synthesis in *Drosophila* motor and sensory neurons by ∼30 to 60%, depending on the specific mutation, cell type, and ANL labeling time.^19,20^ This inhibition of protein synthesis is attributable to sequestration of tRNA^Gly^ by CMT-mutant GlyRS, resulting in insufficient supply of glycyl-tRNA^Gly^ to the ribosome and ribosome stalling on glycine codons.^20^ Thus, we generated *Drosophila* lines that co-express G240R mutant GlyRS and mCD8::GFP in motor neurons (OK371-GAL4), and exposed larvae to 4 mM βES for 48 h. *In vivo* THRONCAT revealed that in motor neurons (identified as mCD8::GFP expressing cells in the VNC) the labeling intensity was reduced by ∼40% upon expression of GlyRS_G240R, as compared to motor neurons expressing mCD8::GFP alone (Fig. 5f,g). This result demonstrates that THRONCAT can be used to quantify and detect cell-type-specific changes in protein synthesis rates in *Drosophila* when combined with fluorescent cell markers.

## Discussion

In this work we have used non-canonical amino acid βES to establish a new metabolic protein labeling technique, THRONCAT. βES is incorporated biosynthetically into nascent proteins in the position of threonine, presumably through ThrRS catalyzed aminoacylation of tRNA^Thr^. We showed that βES competes efficiently with threonine for incorporation into NSPs, suggesting a similar specificity constant for aminoacylation by ThrRS as observed for threonine. Also, we note that threonine residues, compared to methionine, are more abundant in the human proteome (5.2% vs 2.5%) and are often solvent exposed in proteins.^21^ Together, these factors likely contribute to the strong NSP labeling observed when using βES.

Methionine starvation, essential to ensure efficient incorporation of methionine analogs in proteins using BONCAT, has been reported to reduce histone synthesis, biomolecule methylation and to impede cell cycle progression.^22,23^ Indeed, we also observed a reduced proliferation rate of HeLa cells under methionine starvation conditions already at low HPG concentrations. The rapid and strong proteomic incorporation of βES in cells grown in complete medium obviates the need for amino acid starvation. We expect THRONCAT to be especially suited for NSP labeling in fastidious cell types, such as primary cells, that may need specialized media to support their growth. Moreover, THRONCAT enables facile NSP labeling in wild-type strains of bacteria, thereby precluding the need for auxotrophic strains and increasing the scope of bacteria amenable to metabolic protein labeling. Notably, using THRONCAT in *E. coli* we did not observe the impaired bacterial growth that was previously observed with BONCAT.^24^

Although our proteomic experiments in HeLa cells show that THRONCAT provides thorough NSP enrichment, a fraction of proteins was enriched exclusively by BONCAT. In this context, we note that 2.1% of protein sequences in the human proteome do not contain threonine residues and are therefore not amenable to THRONCAT labeling. Conversely, 2.7% of protein sequences does not contain methionine and 9.1% of proteins contain a single N-terminal methionine residue that may be post-translationally removed (UNIPROT human reference proteome UP000005640 analyzed with python script; Supplementary Data 2).

In experiments where exhaustive identification of NSPs is crucial, a combination of THRONCAT and BONCAT may be used to increase proteomic coverage. We believe THRONCAT will find wide application for analysis of proteomic changes such as cellular activation and differentiation, by facilitating simple and fast pulse labeling of NSPs. Although label-free quantification (LFQ) provides simple quantification of NSPs,^25^ we suggest that THRONCAT is compatible with stable isotope labeling by amino acids in culture (SILAC) or tandem mass tags (TMT) for more accurate quantification of protein expression.

Using a *Drosophila* model of CMT peripheral neuropathy, we showed that THRONCAT enables *in vivo* quantification of protein synthesis rates. Although βES incorporation is not inherently cell-type specific, a combination of THRONCAT and GFP-expression in a cell type of interest enabled cell-type specific visualization of protein synthesis. We envision that a similar approach in (co)-cultured cells or organisms may enable cell-specific protein synthesis visualization and quantification.

In summary, we introduced THRONCAT, a novel non-canonical amino acid tagging method based on bioorthogonal threonine analog βES, which enables efficient and non-toxic labeling of newly synthesized proteins in complete growth media within minutes. We foresee that the efficient incorporation of βES in NSPs without the need to deplete threonine and combined with the ease of use, creates unprecedented opportunities to examine cell responses and mechanisms in models that are currently challenging to study.

## Methods

### Reagents and chemical synthesis

All commercial chemicals and solvents were purchased from Sigma Aldrich, Fluorochem, TCI or Fisher Scientific and used without further purification. The identity and purity of each product was characterized by ^1^H NMR, ^13^C NMR, HRMS, TLC, and in the case of chiral products, optical rotation. Absolute stereochemistry of chiral products was determined by Mosher ester analysis.^26^ Stock solutions of amino acid analogs were prepared in Milli-Q water at 200 mM concentrations. Metabolic labeling reporter β-ethynylserine (βES) is available from the authors on request. For detailed synthetic procedures, see Supplementary Note 1.

### Cell culture media

Dulbecco’s modified Eagle’s medium (DMEM, Thermo Fisher, Cat. No. 41965062) was used as complete medium used for metabolic labeling experiments with βES or HPG in HeLa cells, while SILAC RPMI 1640 (RPMI 1640, cat. No: A2494201, supplemented with 300 mg/L *L*-Glutamine, 200 mg/L *L-*Arginine, 2.00 g/L *D*-Glucose, was metabolic labeling in Ramos B cells. Media were supplemented with 10% FCS and 100 units/mL penicillin and 100 μg/mL streptomycin before use. Methionine-free medium and threonine-free medium were custom made in-house. Their formulations were based on DMEM and are outlined in Supplementary Table S2. Methionine-free and threonine-free media were sterile filtered and supplemented with 10% dialyzed FBS (Fisher Scientific, art. No. 15605639) and 100 units/mL penicillin and 100 μg/mL streptomycin before use.

### Cell lines and culture conditions

HeLa cells were cultured in Dulbecco’s modified Eagle’s medium (DMEM, Cat No. 41965062) supplemented with 10% fetal calf serum (FCS), 100 units/mL penicillin and 100 μg/mL streptomycin. Ramos 3F3 cells^27^ were cultured in RPMI 1640 medium (Fisher Scientific, Cat. No.: 11544526 supplemented with 10% FCS, 100 units/mL penicillin and 100 μg/mL streptomycin. The cells were grown at 37 °C in humidified atmosphere of 5% CO_2_ and were used within 15 passages for experiments.

### Amino acid analog labeling in B834 and BL21 *E. Coli*

10 mL of Lysogeny broth (LB) was inoculated from a glycerol stock of *E. Coli* B834 or BL21 and grown overnight in an orbital shaker at 37 °C. The cultures were diluted 10 × with LB medium and the bacteria were grown at 37 °C until OD_600_ was 0.6 – 0.8. Then, bacteria were collected by centrifugation (5,000 *g*, 3 min). *E*. coli B834 (a gift from S. van Kasteren, Leiden University) were incubated in SelenoMet medium (Molecular Dimensions) at 37 °C for 30 min to deplete intracellular methionine stores. Then, the bacteria were collected by centrifugation (5,000 *g*, 3 min) and were incubated at 37°C for 1 h in 1 mL of SelenoMet medium supplemented with either 4 mM HPG or 4 mM βES and 4 mM methionine. *E. coli* BL21 were not starved and incubated directly at 37 °C for 1 h in 1 mL of LB supplemented with either 4 mM βES or 4 mM HPG. *E. coli* BL21 negative controls were incubated with 1 mL of either of 4 mM βES and 34 μg/mL chloramphenicol (CAP) in LB medium or 4 mM βES and 200 mM threonine in LB medium. To asses βES incorporation over time, *E. coli* BL21 were incubated with 1 mL of 4 mM βES in LB medium for the indicated duration, after which βES incorporation was stopped by addition of 34 μg/mL chloramphenicol. After incubation with amino acid analogs, both B834 and BL21 *E. coli* were collected by centrifugation, washed once with PBS and resuspended in 100 μL of lysis buffer (50 mM HEPES, pH 7.4, 150 mM NaCl, 1% Triton-X-100, 1 × EDTA-free protease inhibitor cocktail). Bacteria were lysed using a tip sonicator (MSE, Soniprep 150) for 3 × 10 seconds on ice. Lysate was cleared by centrifugation (13,000 *g*, 30 min) at 4 °C, after which the supernatant was transferred to a new tube and the protein concentration was determined by BCA assay (Thermo Fisher). Lysates were flash frozen and stored at – 80 °C until further use.

### Metabolic labeling of HeLa cells for in-gel analysis

HeLa cells were seeded at a density of 3 × 10^5^ cells per well in a 6 well plate and grown for 1 d in complete medium (DMEM). Then, for labeling experiments performed in threonine-free or methionine-free medium, HeLa cells were starved of intracellular threonine or methionine by incubating at 37 °C in threonine-free or methionine-free medium for 30 min. Then, cells were incubated at 37 °C for 1 h with 4 mM βES or 4 mM HPG in threonine-free or methionine-free medium, respectively. HeLa cells labeled in complete medium (DMEM) were not starved, but were directly incubated with 4 mM βES or 4 mM HPG in complete medium (DMEM) at 37 °C for 1 h. Following incubation, cells were washed with PBS, trypsinized and washed with PBS again. Then, cells were resuspended in 100 μL of lysis buffer (50 mM HEPES, pH 7.4, 150 mM NaCl, 1% Triton-X-100, 1 × EDTA-free protease inhibitor cocktail (Roche)) and lysed using a tip sonicator (MSE, Soniprep 150) for 3 × 10 seconds on ice. Lysate was cleared by centrifugation (13,000 *g*, 30 min) at 4 °C, after which the supernatant was transferred to a new tube and the protein concentration was determined by BCA assay (Thermo Fisher). Lysates were flash frozen and stored at – 80 °C until further use.

### In-gel analysis of *E. coli* and HeLa lysates

100 μg of protein lysate was transferred to a new tube and was diluted to 90 μL total volume using lysis buffer (50 mM HEPES, pH 7.4, 150 mM NaCl, 1% Triton-X-100, 1 × EDTA-free protease inhibitor cocktail). Then 10 μL of a 10x concentrated CuAAC reagent mixture, consisting of 10 mM CuSO_4_ · 5 H_2_O, 20 mM Tris-hydroxypropyltriazolylmethylamine (THPTA), 10 μM Sulfo-Cy5-azide (Jena Bioscience) and 100 mM sodium ascorbate, was added and the reaction mixture was agitated at 37 °C in an orbital shaker at 550 rpm for 30 minutes. Then, protein lysate was precipitated by the addition of 400 μL of ice-cold acetone and protein pellet was collected by centrifugation (5,000 *g*, 20 min) at 4 °C. Protein pellet was allowed to dry on air and was dissolved in 1 × Laemmli sample buffer (Bio-rad). Samples were heated at 70 °C for 10 min and 20 μg of protein for each sample was loaded onto a 1.5 mm 12% polyacrylamide gel. Gels were run at 80V for the stacking gel and 120V for the running gel. All samples were subjected to identical CuAAC labeling conditions, including all negative controls. In-gel fluorescence was measured with a Typhoon™ 5 gel scanner (Cytiva) using the 635 nm laser and the 670BP30 filter. Total protein amount was visualized by silver stain and imaged on a GelDoc XRS+ (Bio-rad).

### Metabolic labeling of HeLa cells for flow cytometry

HeLa cells were seeded at a density of 60,000 cells per well in a 48 well plate and grown for 1 d in complete medium (DMEM). Then, for labeling experiments performed in threonine-free or methionine-free medium, HeLa cells were starved of intracellular threonine or methionine by incubating at 37 °C in threonine-free or methionine-free medium for 30 min. Then, cells were incubated at 37 °C for 1 h with various concentrations of βES or HPG in threonine-free or methionine-free medium, respectively. Negative controls for βES labeling were supplemented with either 100 μM cycloheximide or an excess (1 mM) of threonine. HeLa cells labeled in complete medium (DMEM) were not starved, but were directly incubated with various concentrations of βES or HPG at 37 °C for 1 h. After incubation with analog, all cells were chased by incubation in complete medium (DMEM) for 1 h at 37 °C. For time course analysis of incorporation, HeLa cells were incubated with 4 mM βES or 4 mM HPG for the indicated duration and cells were rinsed once with complete medium (DMEM) and labeling was stopped abruptly by addition of 600 μM cycloheximide. Following chase or cycloheximide treatment, cells were washed twice with cold PBS, trypsinized and fixed with 4% paraformaldehyde for 10 min. Cells were washed twice with PBS and stored at 4 °C until further use.

### Metabolic labeling of HeLa cells for microscopy

An 8-well chamber slide (Ibidi) was coated by treatment with 0.1% gelatin for 30 min at room temperature. The gelatin solution was removed and HeLa cells were seeded at a density of 50,000 cells per well and grown in complete medium (DMEM) at 37 °C for 1 d. Then, cells were incubated for 10 min in complete medium (DMEM) supplemented with either 4 mM βES or 4 mM HPG.

Control cells were left untreated. Amino acid labeling was stopped by quickly aspirating the medium and replacing it by complete medium (DMEM) containing 600 μM cycloheximide. Then, cells were washed twice with cold PBS and fixed with 4% paraformaldehyde for 10 min. washed twice with PBS and stored at 4 °C until further use.

### Competition assay of analogs versus natural amino acids

HeLa cells were seeded at a density of 60,000 cells per well in a 48 well plate and grown for 1 d in complete medium (DMEM). Cells were starved of intracellular threonine or methionine by incubating at 37 °C in threonine-free or methionine-free medium for 1 h. Then, cells were incubated 37 °C for 1 h with 50 μM βES and various ratios of threonine or 50 μM HPG and various ratios of methionine in threonine-free or methionine-free medium, respectively. After incubation, cells were washed twice with cold PBS, fixed with 4% paraformaldehyde for 10 minutes, washed twice with PBS and stored at 4 °C until further use. Incorporated analogs were conjugated to Cy5-azide and quantified by flow cytometry.

### Cy5 conjugation to HeLa proteome via azide-alkyne click chemistry

Metabolic labeling for flow cytometry and microscopy was performed as previously described. Then, cells were permeabilized by incubation in 200 μL of 0.3% saponin in PBS for 10 min. Next, cells were incubated in 200 μL of a blocking buffer (0.1% saponin, 3% BSA in PBS) for 30 min. Copper-catalyzed azide-alkyne click (CuAAC) reaction was perfomed for 30 min at 37 °C with 100 μL of a CuAAC reaction mixture (4 mM CuSO4 · 5 H2O, 200 μM THPTA, 0.5 μM sulfo-Cy5-azide and 40 mM sodium ascorbate, 0.1% saponin in PBS). Then, cells were washed 4 × with 200 μL blocking buffer (0.1% saponin, 3% BSA in PBS) and resuspended in 3% BSA in PBS for flow cytometry or resuspended in PBS for widefield microscopy. All replicates were subjected to identical CuAAC labeling conditions, including all negative controls.

### Cell viability assay

HeLa cells were seeded at a density of 10,000 cells per well into a 96 well plate and grown for 1 d in complete medium (DMEM). Cells were then incubated for the indicated amount of time various concentrations of βES in complete medium (DMEM) or various concentrations of HPG in methionine-free medium. Cells grown in complete medium (DMEM) without analog were used as a control. Following incubation, cells were trypsinized and resuspended in BD FACS flow buffer supplemented with 3 μM propidium iodide (Sigma). Propidium iodide exclusion was analyzed by flow cytometry.

### CellTrace Violet assay

1 × 10^6^ HeLa cells were suspended in 10 mL of a 1 μM CellTrace violet solution in PBS and incubated at 37 °C for 20 min. Excess CellTrace dye was quenched by addition of 10 mL of complete medium (DMEM). Cells were split resuspended in either complete medium (DMEM) without analog, complete medium (DMEM) supplemented with various concentrations of βES or methionine-free medium supplemented with various concentrations of HPG. Cells were seeded at a density of 1 × 10^5^ cells per well in a 12 well plate and incubated at 37 °C for 24 h. After trypsinization, cell proliferation was assessed by quantifying residual mean CellTrace fluorescence by flow cytometry. CellTrace-labeled HeLa cells at *t* = 0 were measured to establish baseline fluorescence.

### Flow cytometry

Flow cytometry was performed on a FACSVerse™ (BD Biosciences) flow cytometer. Cy5 was detected using the 633 nm laser line and the 660/10 filter, CellTrace violet using the 405 nm laser line and the 448/45 filter and propidium iodide using the 488 nm laser and the 700/54 filter.

### Widefield microscopy

HeLa cells were metabolically labeled and subjected to CuAAC with Cy5-azide as previously described. Cells were imaged on a DMi8 widefield microscope (Leica). Images were processed and analyzed in ImageJ/Fiji (National Institutes of Health). Image contrast was enhanced to improve signal visibility by changing the maximum displayed values. Maximum intensity projection was used in all images.

### THRONCAT and BONCAT in HeLa cells for proteomics

HeLa cells were seeded at a density of 4 × 10^6^ cells per T-75 flask and grown at 37 °C for 1 d. For HPG labeling, cells were incubated in methionine-free medium at 37 °C for 30 min to deplete intracellular methionine. Then, cells were incubated for 5 h at 37 °C with 4 mM HPG in methionine-free medium. HeLa cells labeled with βES were not starved, but instead incubated directly in either 1 mM βES or 4 mM βES in complete medium (DMEM) for 5 h at 37 °C. Control cells were incubated for 5 h at 37 °C in complete medium (DMEM) without analog. After incubation, cells were washed twice in PBS, trypsinized and collected by centrifugation. Cells were washed again in PBS and stored at – 80 °C until further use.

### THRONCAT labeling in Ramos cells for proteomics

1 × 10^7^ Ramos 3F3 cells were collected for each replicate by centrifugation. For steady state replicates, cells were resuspended in 5 mL of SILAC RPMI 1640 (RPMI 1640, cat. No: A2494201, supplemented with 300 mg/L *L*-Glutamine, 200 mg/L *L-*Arginine, 2.00 g/L *D*-Glucose and 10% dialyzed FBS) with 146 mg/L ^13^C_6_,^*15*^*N*_*2*_*-*L-lysine (Lys_8_, Silantes, Prod. No 211604102) and 1 mM βES and grown at 37 °C for 1 h. Simultaneously, for anti-IgG stimulated replicates, cells were resuspended in RPMI 1640 supplemented with 10 μg/mL of anti IgG (Invitrogen, Cat. No. 31143). For anti-IgG stimulated replicates that were pulse labeled between 1-2 h or 3-4h, cells were resuspended in RPMI 1640 with 10 μg/mL anti-IgG and grown at 37 °C for 1 or 3 h, followed by centrifugation and incubation at 37 °C in SILAC RPMI 1640 with 10 μg/mL anti-IgG, 146 mg/L Lys_8_ and 1 mM βES. For anti-IgG stimulated replicates that were pulse labeled between 0-1 h, cells were resuspended in SILAC RPMI 1640 with 10 μg/mL anti-IgG, 146 mg/L Lys_8_ and 1 mM βES and grown at 37 °C for 1 h. For all replicates, pulse labeling was stopped by the addition of 600 μM cycloheximide and cells were collected by centrifugation. Cells were washed once with PBS and cell pellets were flash frozen and stored at – 80 °C until further use.

### Enrichment of nascent protein and on-bead digestion

HeLa or Ramos cells were metabolically pulse-labeled as previously described. Then, cell pellets were resuspended in SDS lysis buffer (4% SDS, 1 mM DTT, 100 mM Tris-HCl pH 7.5) and were incubated for 5 minutes at 95 °C. Samples were sonicated until homogeneous using alternating cycles of 30 sec on/30 sec off on the highest setting using a BioRuptor Pico (Diagenode) and spun down at 16,000 × *g* for 5 min. Supernatant was transferred to a new tube and protein concentrations were determined using Pierce BCA protein assay (ThermoFisher, 23225). Samples were stored at -80 °C until further use. Then, 400 μg and 200 μg whole cell lysate in 250 uL PBS was used as input material for HeLa cells and Ramos cells, respectively. CuAAC reaction was set up following the instructions from the Click-iT Protein Enrichment Kit (Invitrogen, C10416), but with the provided beads substituted for azide agarose beads (Sigma Aldrich, 900957). 100 μL bead slurry was used per reaction, which was washed with 1 mL H_2_O, after which the cell lysate, 250 μL urea buffer, 500 μL 2x catalyst solution were added and incubated for 16 -20 hours at room temperature with end-over-end rotation, after which the beads were washed with 1 mL of H_2_O. Then, beads were incubated with 10 mM DTT in 0.5 mL SDS buffer (provided by the kit) at 70 °C for 15 min while shaking. The beads were spun down, the supernatant was aspirated and the beads were incubated with 50 mM iodoacetamide in 0.5 mL SDS buffer for 30 min at room temperature while shaking. The beads were transferred to spin columns (provided by the kit) and washed with 20 mL of SDS buffer, 20 mL of 8 M urea in 100 mM Tris, pH 8, 20 mL of 20% isopropanol, 20 mL of 20% acetonitrile and 5 mL of PBS. The bound proteins were digested by resuspending the beads in 200 μL freshly prepared digestion buffer (2M Urea, 100 mM Tris-HCl pH 8, 100 mM DTT) with 0.5 ug trypsin (Promega, V5111) and overnight incubation at room temperature while shaking. The digest was collected and acidified with 10 μL 10% CF_3_COOH, after which the peptides were desalted and stored on StageTips.^28^

### LC-MS/MS measurements and data analysis

Peptide samples were eluted from StageTips with elution buffer (80% acetonitrile, 0.1% formic acid in H2O), reduced to 10% of the original volume by vacuum concentration and diluted in 0.1% formic acid. Peptides were separated using an Easy-nLC 1000 liquid chromatography system (ThermoScientific) with a 44 minute acetonitrile gradient (7-30%), followed by washes at 60% and 95% acetonitrile for a total of 60 minutes data collection. Mass spectrometry was performed on an Orbitrap Exploris 480 (ThermoScientific) in data-dependent top-20 mode with dynamic exclusion set at 45 seconds. Protein identification and quantification was done in MaxQuant v1.6.0.1^29^ with match-between-runs, iBAQ and label-free quantification enabled. Methionine-to-HPG (−22.0702 Da; HeLa experiment) and threonine-to-βES (+9.984 Da; HeLa and Ramos experiment) were added to the default variable modifications. For proteomic analysis of Ramos NSPs, heavy arginine (Arg_6_) and Lys_8_ were set as labels to exclusively use Lys_8_-containing peptides for protein identification and quantification. The MS/MS spectra were searched against a human UniProt database downloaded in June 2017. Common contaminants and decoy database hits were removed from the resulting MaxQuant output files and alias gene names were replaced with official gene symbols using the Limma package.^30^ If this resulted in duplicated entries, only the entry with the highest Andromeda Score was retained. Only proteins that were detected in all replicates of a condition were marked as identified protein. For HeLa proteomic analysis, all proteins that were detected in replicates of the untreated negative control were considered a-specific binders and were removed for downstream analysis for all conditions. Average threonine-or methionine content of protein fractions was determined by a Python script (Supplementary Data 2). Uniprot reference proteome (UP000005640) was used as complete human proteome to analyze average proteomic threonine and methionine content.

Differentially enriched protein analysis was performed using the DEP package.^31^ All proteins that were detected in all replicates of at least one condition were considered for downstream analysis. Imputation of missing values was performed using the MinProb method with the default settings. Imputation was performed 1000x and adjusted p-values and fold changes in LFQ intensities were calculated for each round. All proteins that showed an adjusted p-value < 0.05 and a fold change > 1.5 in more than 80% of the iterations were considered to be significantly differentially expressed.

The mass spectrometry proteomics data have been deposited to the ProteomeXchange Consortium (http://proteomecentral.proteomexchange.org) via the PRIDE partner repository^32^ with the dataset identifier PXD032368.

### Drosophila genetics

Flies were housed in a temperature-controlled incubator with 12:12 h on/off light cycle at 25°C. OK371-GAL4, UAS-mCD8::GFP; UAS-mCD8::GFP was kindly provided by Dr. M. Freeman (Vollum Institute, Oregon Health & Science University, USA). The UAS-hGlyRS_G240R line was previously described.^19^ UAS-hGlyRS_G240R or w1118 (control) flies were crossed with OK371-GAL4, UAS-mCD8::GFP; UAS-mCD8::GFP flies and larval offspring was used in experiments.

### THRONCAT in *Drosophila melanogaster*

For *in vivo* THRONCAT, previously described FUNCAT procedures^18–20,33^ were adapted. 8 h egg collections were performed and animals were raised on Jazz-Mix Drosophila medium (Fisher Scientific) at 25 °C. 72 h after egg laying (AEL), larvae were transferred to βES or HPG-containing medium. The standard βES or HPG concentration used was 4 mM, except for the βES dosage-titration experiment (Fig. 5e). The standard exposure time to non-canonical amino acid was 48 h, except in the experiment shown in Fig. 5f, in which larvae were exposed to βES for different time frames (2 h, 4 h, 8 h, 16 h, or 48 h). Larval central nervous systems (CNSs) were dissected in ice-cold HL3 solution and fixed in 4% paraformaldehyde for 30 min at room temperature. After fixation, the CNSs were washed 3 × 15 min with PBST (1x PBS pH 7.2 containing 0.2% Triton X-100) and 3 × 15 min with PBS pH 7.2. Metabolically labeled NSPs were conjugated to tetramethylrhodamine 5-carboxamido-(6-azidohexanyl) (TAMRA-N_3_, Invitrogen, T10182) via CuAAC. The CuAAC reaction mix was assembled in a defined sequence of steps. THPTA ligand (200 μM), TAMRA-N_3_ (2 μM), CuSO_4_ solution (4 mM) and sodium ascorbate solution (40 mM) were added to PBS pH 7.2. After each addition the solution was mixed thoroughly for 10 sec and at the end for 30 sec using a high-speed vortex. Larval CNSs were incubated with 500 μl of CuAAC reaction mix overnight at 4 °C on a rotating platform. The next day, the brains were washed 3 × 15 min with PBS-Tween (1x PBS pH 7.2 containing 1% Tween-20) and 3 × 15 min with PBST. Finally, larval brains were mounted in VectaShield mounting medium (Biozol, VEC-H-1000-CE) and stored at 4 °C until imaging using a Leica SP8 laser scanning confocal microscope. For image acquisition, identical confocal settings were used for all samples of a given experiment. Fluorescence intensities of the TAMRA signal were quantified using ImageJ/FIJI software (National Institutes of Health). Mean intensity of all cells within one motor neuron cluster from each ventral nerve cord were used as single data points for statistical analysis.

### Statistics

For flow cytometry experiments, at least three replicates were measured for each condition (*n* = 3). A minimum number of *n* = 10.000 events were measured (before gating) for each replicate. For experiments with *Drosophila melanogaster*, no statistical methods were used to pre-determine sample sizes, but sample sizes are similar to those reported in previous publications.^19,20^ Whenever possible, data collection and analysis were performed by investigators blinded to the genotype of the experimental animals. Animals or samples were assigned to the various experimental groups based on the concentration and time frame of exposure to βES or HPG, or their genotype. For a given experiment, samples from the different experimental groups were processed in parallel and analyzed in random order. Before analysis, a Robust regression and Outlier removal method (ROUT) was performed to detect all outliers. Normality of all data was analyzed by Shapiro–Wilk, Anderson-Darling and Kolmogorov-Smirnov tests. Subsequent statistical tests were only performed if all assumptions were met. Unpaired t-test was used for comparisons of two groups of normally distributed data with homogeneous variance. Mann-Whitney test was used for comparison of two groups of not normally distributed data. Brown-Forsythe ANOVA with Dunnett’s T3 multiple comparisons test was used for comparison of more than two groups of normally distributed data with non-homogeneous variance. For comparison of more than two groups of not normally distributed data, the Kruskal-Wallis test with Dunn’s multiple comparisons test was used. Statistical analysis was performed using GraphPad Prism v.9.2.0.

## SUPPLEMENTARY INFORMATION

Supporting information describes the chemical synthesis and analysis data of βES, supplementary methods and figures.

### AUTHOR CONTRIBUTIONS

B.J.I and K.B. designed the project. B.J.I performed organic synthesis and THRONCAT experiments in bacteria and HeLa cells. B.J.I. and M.J.v.W performed THRONCAT labeling on Ramos B-cells. J.D. and M.V. performed nascent protein enrichment, proteomics experiments and analysis of proteomics data. E.Sl., N.M.G., E.St., performed and analyzed the data of *Drosophila* experiments. B.J.I and K.B, wrote the main body of the manuscript. E. Sl, N.M.G and E. St wrote the Drosophila results and methods of the manuscript. J.D. and M.V. wrote proteomic methods of the manuscript. B.J.I, E. Sl, N.M.G and E. St designed graphical contents. All authors provided critical advice during the experimental phase and writing of the manuscript. All authors have given permission for publishing of the manuscript.

## ACKNOWLEDGEMENTS

This work is part of a project that has received funding from the European Research Council (ERC) under the European Union’s Horizon 2020 research and innovation program (grant agreement No 802940) and the NWO gravitation program ‘Institute for Chemical Immunology’ (NWO-024.002.009). E.St is supported by an ERC consolidator grant (ERC-2017-COG 770244), and funding from the Radala Foundation, ‘Stichting ALS Nederland’, AFM-Telethon and ARSLA. The Vermeulen lab is part of the Oncode Institute, which is partly funded by the Dutch Cancer Society (KWF).

## References

1. van Bergen, W., Heck, A. J. R. & Baggelaar, M. P. Recent advancements in mass spectrometry–based tools to investigate newly synthesized proteins. Curr. Opin. Chem. Biol. 102074 (2021).

2. Sletten, E. M. & Bertozzi, C. R. Bioorthogonal Chemistry: Fishing for Selectivity in a Sea of Functionality. Angew. Chem. Int. Ed. 48, 6974–6998 (2009).

3. Dieterich, D. C., Link, A. J., Graumann, J., Tirrell, D. A. & Schuman, E. M. Selective identification of newly synthesized proteins in mammalian cells using bioorthogonal noncanonical amino acid tagging (BONCAT). Proc. Natl. Acad. Sci. 103, 9482–9487 (2006).

4. Beatty, K. E. et al. Fluorescence Visualization of Newly Synthesized Proteins in Mammalian Cells. Angew. Chem. Int. Ed. 45, 7364–7367 (2006).

5. Dieterich, D. C. et al. In situ visualization and dynamics of newly synthesized proteins in rat hippocampal neurons. Nat. Neurosci. 13, 897–905 (2010).

6. Liu, J., Xu, Y., Stoleru, D. & Salic, A. Imaging protein synthesis in cells and tissues with an alkyne analog of puromycin. Proc. Natl. Acad. Sci. 109, 413–418 (2012).

7. Kiick, K. L., Saxon, E., Tirrell, D. A. & Bertozzi, C. R. Incorporation of azides into recombinant proteins for chemoselective modification by the Staudinger ligation. Proc. Natl. Acad. Sci. 99, 19–24 (2002).

8. McMurry, J. L. & Chang, M. C. Y. Fluorothreonyl-tRNA deacylase prevents mistranslation in the organofluorine producer Streptomyces cattleya. Proc. Natl. Acad. Sci. 114, 11920–11925 (2017).

9. Minajigi, A., Deng, B. & Francklyn, C. S. Fidelity Escape by the Unnatural Amino Acid β-Hydroxynorvaline: An Efficient Substrate for Escherichia coli Threonyl-tRNA Synthetase with Toxic Effects on Growth. Biochemistry 50, 1101–1109 (2011).

10. Hartman, M. C. T., Josephson, K. & Szostak, J. W. Enzymatic aminoacylation of tRNA with unnatural amino acids. Proc. Natl. Acad. Sci. 103, 4356–4361 (2006).

11. Sanada, M., Miyano, T. & Iwadare, S. β-ethynylserine, an antimetabolite of L-threonine, from Streptomyces cattleya. J. Antibiot. (Tokyo) 39, 304–305 (1986).

12. Potgieter, H. C., Vermeulen, N. M. J., Potgieter, D. J. J. & Strauss, H. F. A toxic amino acid, 2(S)3(R)-2-amino-3-hydroxypent-4-ynoic acid from the fungus Sclerotium rolfsii. Phytochemistry 16, 1757–1759 (1977).

13. Marchand, J. A. et al. Discovery of a pathway for terminal-alkyne amino acid biosynthesis. Nature 567, 420–424 (2019).

14. Santiago, T. et al. FCRLA is a resident endoplasmic reticulum protein that associates with intracellular Igs, IgM, IgG and IgA. Int. Immunol. 23, 43–53 (2011).

15. Irons, E. E. et al. B cells suppress medullary granulopoiesis by an extracellular glycosylation-dependent mechanism. eLife 8, e47328 (2019).

16. Brescia, P. et al. MEF2B Instructs Germinal Center Development and Acts as an Oncogene in B Cell Lymphomagenesis. Cancer Cell 34, 453-465.e9 (2018).

17. Pore, D. & Gupta, N. The Ezrin-Radixin-Moesin Family of Proteins in the Regulation of B-Cell Immune Response. Crit. Rev. Immunol. 35, (2015).

18. Erdmann, I. et al. Cell-selective labelling of proteomes in Drosophila melanogaster. Nat. Commun. 6, 7521 (2015).

19. Niehues, S. et al. Impaired protein translation in Drosophila models for Charcot–Marie–Tooth neuropathy caused by mutant tRNA synthetases. Nat. Commun. 6, 7520 (2015).

20. Zuko, A. et al. tRNA overexpression rescues peripheral neuropathy caused by mutations in tRNA synthetase. Science 373, 1161–1166 (2021).

21. Rose, G. D., Geselowitz, A. R., Lesser, G. J., Lee, R. H. & Zehfus, M. H. Hydrophobicity of Amino Acid Residues in Globular Proteins. Science 229, 834–838 (1985).

22. Hoffman, R. M. & Yano, S. Tumor-Specific S/G2-Phase Cell Cycle Arrest of Cancer Cells by Methionine Restriction. in Methionine Dependence of Cancer and Aging: Methods and Protocols (ed. Hoffman, R. M.) 49–60 (Springer, 2019).

23. Arnaudo, A. M., Link, A. J. & Garcia, B. A. Bioorthogonal Chemistry for the Isolation and Study of Newly Synthesized Histones and Their Modifications. ACS Chem. Biol. 11, 782–791 (2016).

24. van Elsland, D. M. et al. Detection of bioorthogonal groups by correlative light and electron microscopy allows imaging of degraded bacteria in phagocytes. Chem Sci 7, 752–758 (2016).

25. Cox, J. et al. Accurate Proteome-wide Label-free Quantification by Delayed Normalization and Maximal Peptide Ratio Extraction, Termed MaxLFQ. Mol. Cell. Proteomics 13, 2513– 2526 (2014).

26. Hoye, T. R., Jeffrey, C. S. & Shao, F. Mosher ester analysis for the determination of absolute configuration of stereogenic (chiral) carbinol carbons. Nat. Protoc. 2, 2451–2458 (2007).

27. Kissel, T. et al. Antibodies and B cells recognising citrullinated proteins display a broad cross-reactivity towards other post-translational modifications. Ann. Rheum. Dis. 79, 472–480 (2020).

28. Rappsilber, J., Mann, M. & Ishihama, Y. Protocol for micro-purification, enrichment, pre-fractionation and storage of peptides for proteomics using StageTips. Nat. Protoc. 2, 1896– 1906 (2007).

29. Cox, J. & Mann, M. MaxQuant enables high peptide identification rates, individualized p.p.b.-range mass accuracies and proteome-wide protein quantification. Nat. Biotechnol. 26, 1367– 1372 (2008).

30. Ritchie, M. E. et al. limma powers differential expression analyses for RNA-sequencing and microarray studies. Nucleic Acids Res. 43, e47 (2015).

31. Zhang, X. et al. Proteome-wide identification of ubiquitin interactions using UbIA-MS. Nat. Protoc. 13, 530–550 (2018).

32. Perez-Riverol, Y. et al. The PRIDE database and related tools and resources in 2019: improving support for quantification data. Nucleic Acids Res. 47, D442–D450 (2018).

33. Erdmann, I. et al. Cell Type-specific Metabolic Labeling of Proteins with Azidonorleucine in Drosophila. Bio-Protoc. 7, e2397–e2397 (2017).

