## Supplementary Data 2 for "THRONCAT: Efficient metabolic labeling of newly synthesized proteins using a bioorthogonal threonine analog"

# Find Threonine and Methionine content (in %) of complete proteome or of a list of proteins. 
# Written for Python 3
 
from Bio.SeqUtils.ProtParam import ProteinAnalysis
from Bio import SeqIO


import fastaparser
# Standard plotly import
import plotly.graph_objs as go
import chart_studio.plotly
from plotly.offline import iplot, init_notebook_mode
# Using plotly + cufflinks in offline mode
import cufflinks
cufflinks.go_offline(connected=True)
init_notebook_mode(connected=True)
# Block Pandas warning
import warnings
warnings.simplefilter(action='ignore', category=FutureWarning)
warnings.simplefilter(action='ignore', category=DeprecationWarning)

# Import Pandas and Numpy
import pandas as pd
import numpy as np

proteome = []
Sequence_ID = []
First_Residue = []
Protein_name = []

#Fasta parser
#Insert extension of Fasta file containing proteome or list of proteins of interest. Must be in Fasta format. 
with open('Insert file extension here') as fasta_file:
    parser = fastaparser.Reader(fasta_file)
    for seq in parser:
        # seq is a FastaSequence object
        # add protein sequences in FASTA file to a list called 'proteome'
        proteome.append(seq.sequence_as_string())
        # add protein sequence ID's to an empty list called 'Sequence_ID' 
        Sequence_ID.append(seq.id)
        # add protein name to an empty list called 'Protein_name' 
        Protein_name.append(seq.description)
        # puts amino acid sequence of protein in a variable 'AA_Sequence' and checks if first residue is a methionine.
        # If first residue is a methionine, string 'M' is added to a list called 'First_Residue', else 'other' is added to 
        # that list.
        AA_Sequence = seq.sequence_as_string()
        if AA_Sequence.startswith('M') == True:
            First_Residue.append('M')
        else:
            First_Residue.append('other')        
        
# Function that counts threonine and methionine residues in the protein sequences within the target proteome or list of proteins
# function that takes a list of strings as input and counts the instances of 'T' and 'M' for each individual string
# within that list. The 'T' and 'M' counts are stored for every protein in an inner list ámino_acids. The inner lists 
# are stored in an outer list 'T_M_count'. The function returns the 'T_M_count' outer list.
def AA_ResidueCount(proteome):
    # create an empty outer list for the lists of amino acid counts of the various proteins in proteome
    T_M_count = []
    for protein_sequence in proteome:
        # create an empty inner list for the amino acid counts of the various proteins in proteome
        amino_acids = []
        # parse protein sequences using 'proteinanalysis' from Bio module
        # then count the amount of threonines and methionines
        X = ProteinAnalysis(protein_sequence)
        T=X.count_amino_acids()['T']
        M=X.count_amino_acids()['M']
        # add the amino acids counts to the empty list 'amino_acids'
        amino_acids.append(T)
        amino_acids.append(M)
        # add the list that contains an amino acid count for an individual protein to the outer list 'T_M_count' that holds
        # the inner lists for the whole proteome
        T_M_count.append(amino_acids)
    return T_M_count

def AA_Protein_Fraction(proteome):
    fraction_count = []
    for protein_sequence in proteome:
        # create an empty inner list for the amino acid fractions of the various proteins in proteome
        amino_acids_fractions = []
        # parse protein sequences using 'proteinanalysis' from Bio module
        # then determine the fraction of threonines and methionines
        X = ProteinAnalysis(protein_sequence)
        T_frac="%0.3f" % X.get_amino_acids_percent()['T']
        M_frac="%0.3f" % X.get_amino_acids_percent()['M']  
        # convert fractions from strings into floats
        T_float = float(T_frac)
        M_float = float(M_frac)
        # add the amino acids fractions to the empty list 'amino_acids'
        amino_acids_fractions.append(T_float)
        amino_acids_fractions.append(M_float)
        # add the list that contains an amino acid fractions for an individual protein to the outer list 'fraction_count' that holds
        # the inner lists for the whole proteome
        fraction_count.append(amino_acids_fractions)
    return fraction_count


# Puts residue counts for threonine and methionine in a Pandas dataframe called 'df1'.
# Inserts a column which contains information on the identity of the first residue 
df1 = pd.DataFrame(AA_ResidueCount(proteome), columns=['Threonine', 'Methionine'])
df1.insert(2, "Sequence ID", Sequence_ID, True)
df1.insert(2, "First Residue", First_Residue, True)
df1.insert(2, "Protein Name", Protein_name, True)


# Puts Threonine and Methionine fractions of the proteins in a Pandas dataframe called 'df2'
df2 = pd.DataFrame(AA_Protein_Fraction(proteome), index=Sequence_ID, columns=['Threonine', 'Methionine'])
# Calculates the average of the individual proteins' Threonine fractions, creating the average Threonine content for the
# complete proteome. Multiplies the fraction with 100 to create percentage, then rounds of percentage to 3 decimal points.
fraction_dec_T=df2['Threonine'].mean()
fraction_perc_T = round(fraction_dec_T *100, 3)
#Does the same for Methionine. 
fraction_dec_M=df2['Methionine'].mean()
fraction_perc_M = round(fraction_dec_M *100, 3)

print('Threonine fraction of proteome:',fraction_perc_T, '%')
print()
print('Methionine fraction of proteome:',fraction_perc_M, '%')
print()
print()
print()


# Finds all proteins that contain an X amount of threonines/methionines and stores them in dataframe 3 and 4 respectively.
# the amount of threonine/methionines you are looking for (e.g., 0 or 3 per protein) is represented by the variable 
# 'amino_acid_query'. 
amino_acid_query = 1
df3 = (df1.loc[df1['Threonine'] == amino_acid_query])
print("Number of proteins that contain", amino_acid_query, "threonine residue(s):")
print(len(df3))
print()
df4 = (df1.loc[df1['Methionine'] == amino_acid_query])
print("Number of proteins that contain", amino_acid_query, "methionine residue(s):")
print(len(df4))
print()
print()
print()
df5 = df4.loc[df4["First Residue"] == "M"] 
print("Number of proteins that contain", amino_acid_query, "methionine residue(s), with methionine being starting (N-terminal) residue:")
print(len(df5))
