## Supplementary Figures for "THRONCAT: Efficient metabolic labeling of newly synthesized proteins using a bioorthogonal threonine analog"

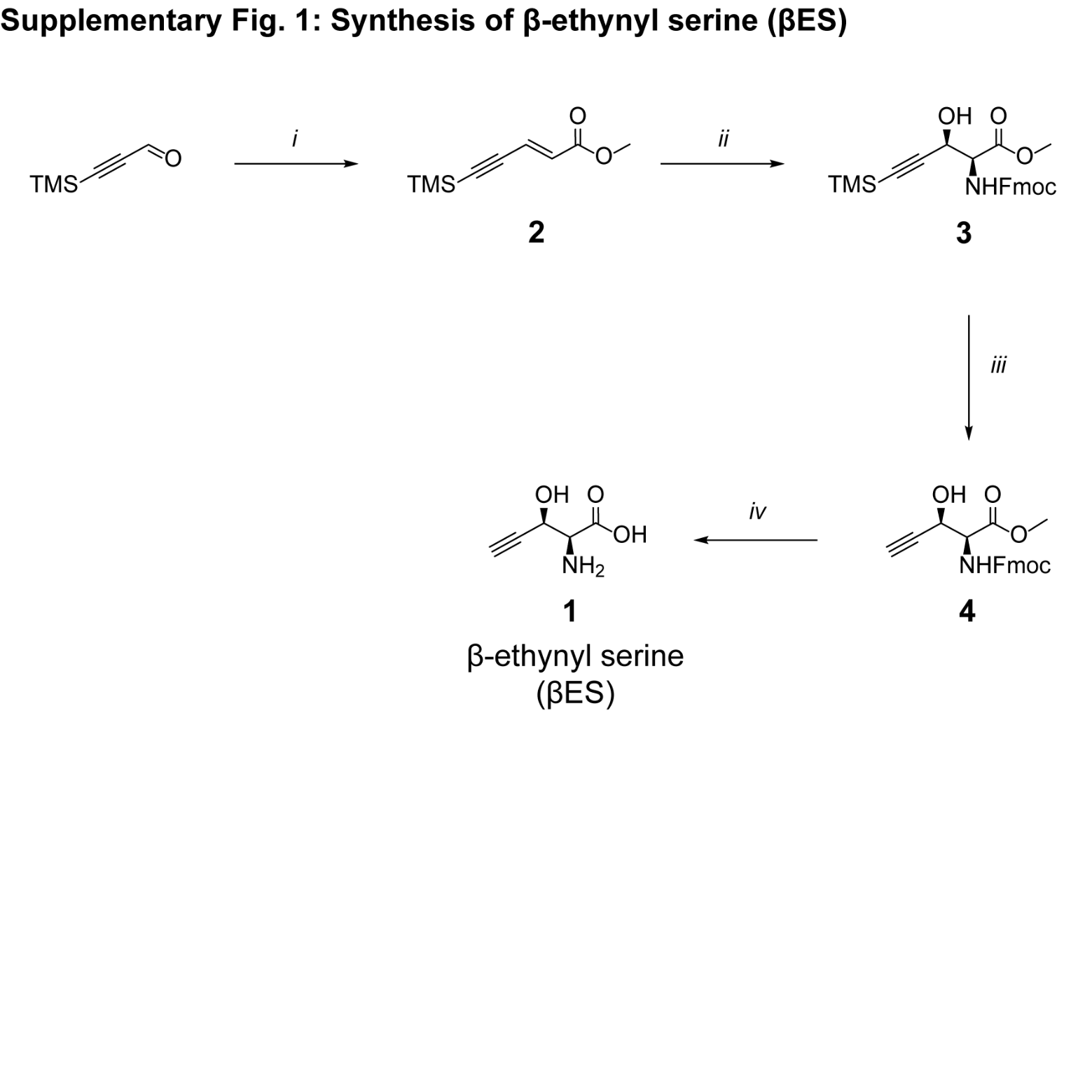

**Supplementary Fig. 1:** **Synthesis of (2S,3R)-2-amino-3-hydroxypent-4-ynoic acid (β-ethynylserine, 1).** Reagents and yields: i) methyl (triphenylphosphoranylidene)acetate, THF, 95%; ii) (DHQD)2AQN, K2OsO4 · 2 H2O, NaOH, 5, H2O/n-PrOH, 31%; iii) TBAF, DCM, 82%; iv) LiOH, H2O/MeCN, 93% (86% ee).

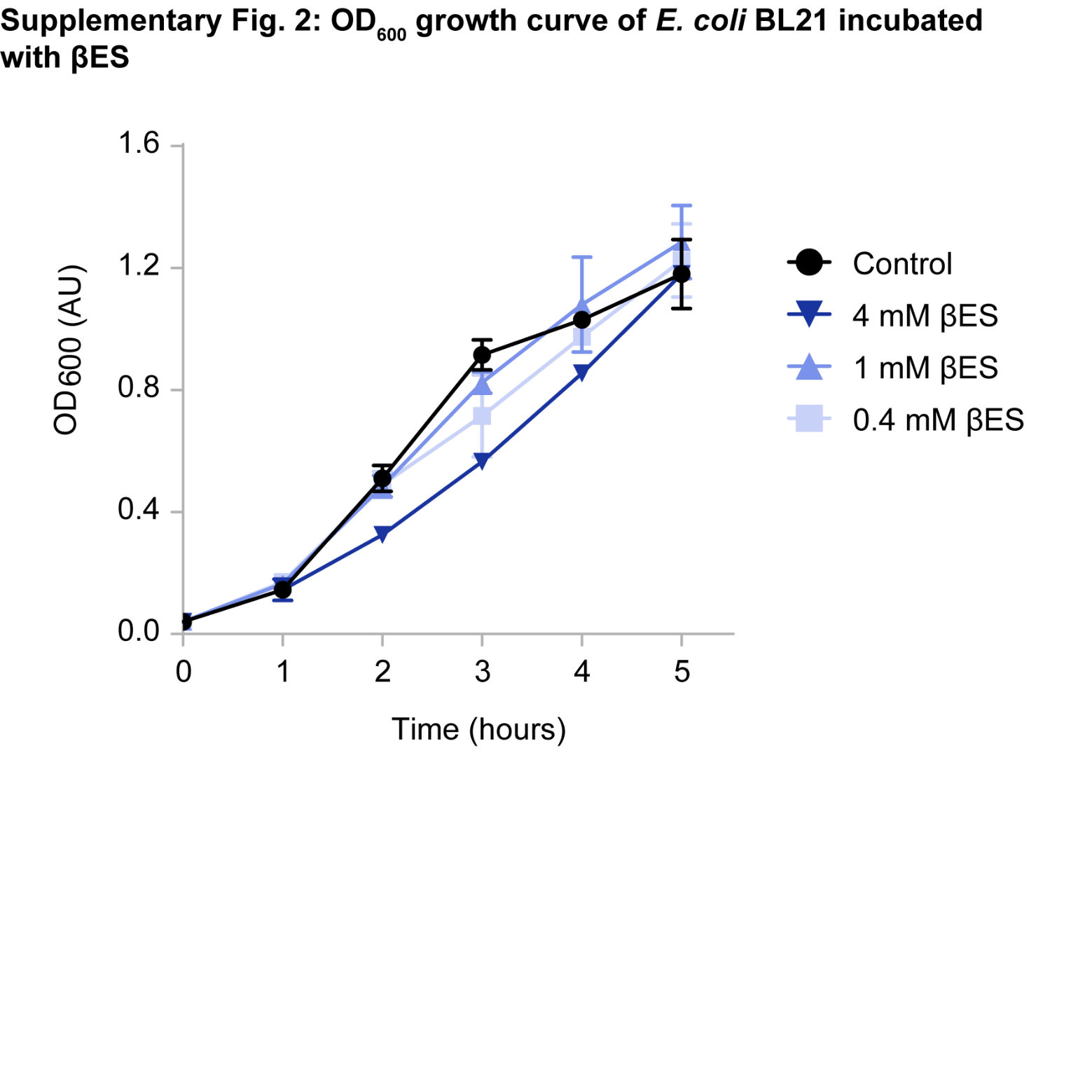

**Supplementary Fig. 2: Growth curve of E. coli BL21 incubated with various concentrations of βES**. E. coli were incubated with 0.4, 1 or 4 mM βES in LB medium and bacterial growth was monitored by OD_600_ absorbance. E. coli grown in LB medium without βES were included as a control. OD_600_, optical density at 600 nm; AU, arbitrary units. Error bars represent s.d., Sample size is n = 2.

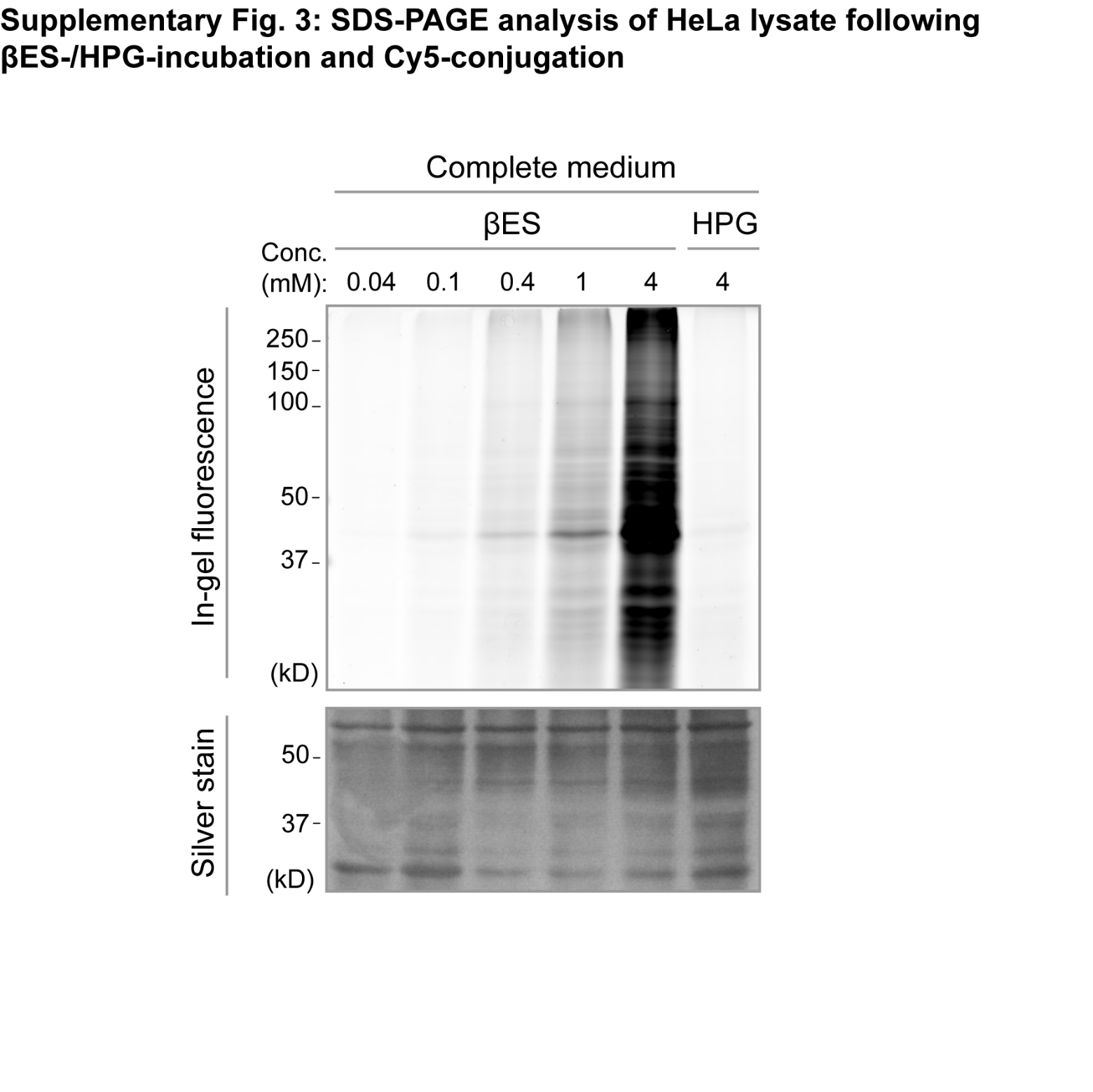

**Supplementary Fig. 3: In-gel visualization of βES or HPG incorporation into the HeLa proteome**. *E. coli* were incubated for 1 h with indicated concentrations of βES or HPG in LB medium. *E. coli* lysate was conjugated to Cy5-azide and visualized by in-gel fluorescence after SDS-PAGE separation. Silver stain panel shows total protein in lysates.

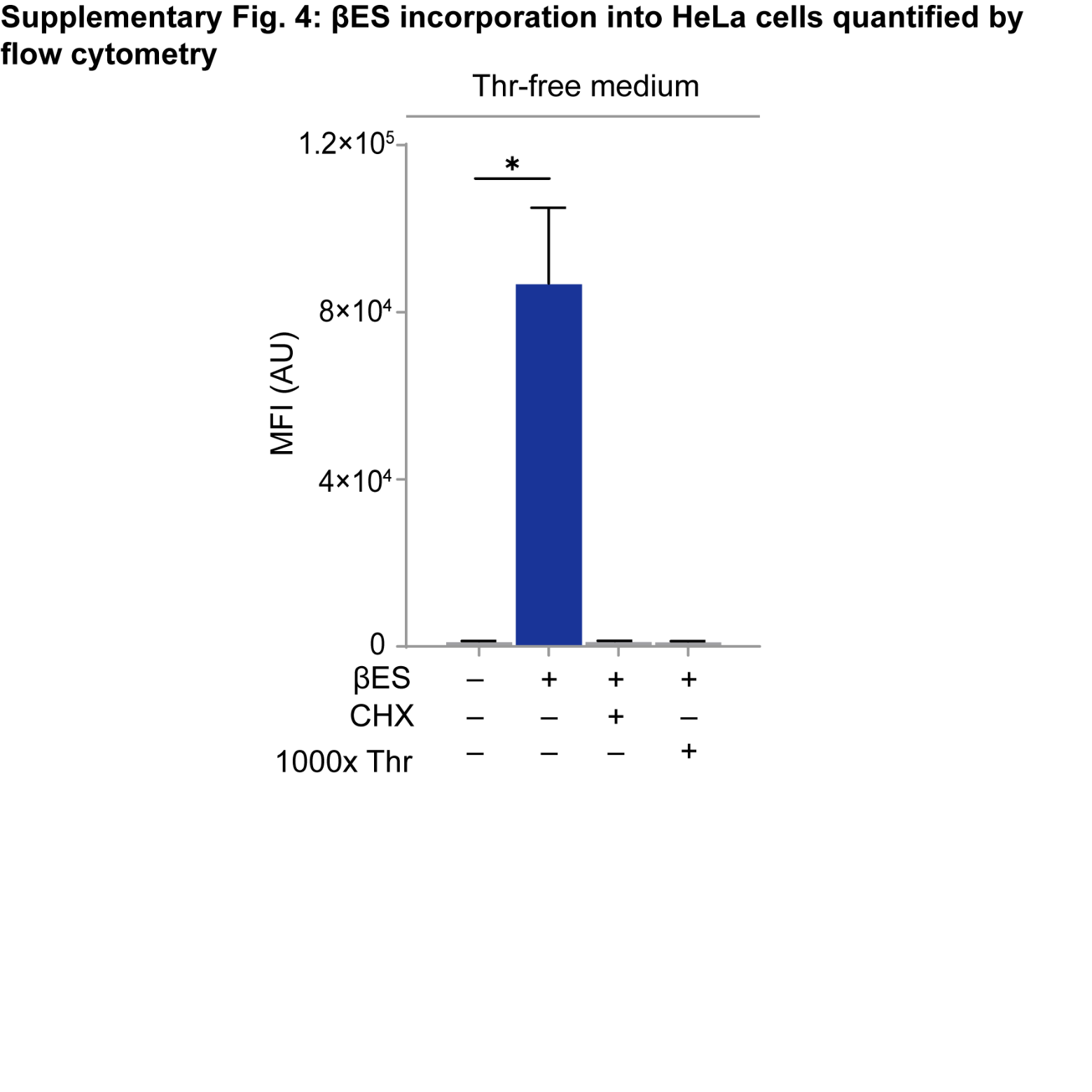

**Supplementary Fig. 4: Flow cytometry quantification of βES incorporation into HeLa proteome**. HeLa cells were starved for 1 h in threonine-free medium and incubated for 1 h with 1 μM βES in threonine-free medium. Control cells were left untreated or treated with 1 μM βES and protein synthesis inhibitor cycloheximide or, 1 μM βES and an excess of threonine. Incorporated βES was conjugated to Cy5-azide for quantification. MFI, Mean fluorescence intensity; AU, arbitrary units; CHX, cycloheximide; Thr, threonine. **P =* 0.0146 (95% confidence interval); determined by an unpaired two-tailed Students t test. Error bars represent s.d. Sample size is *n* = 3.

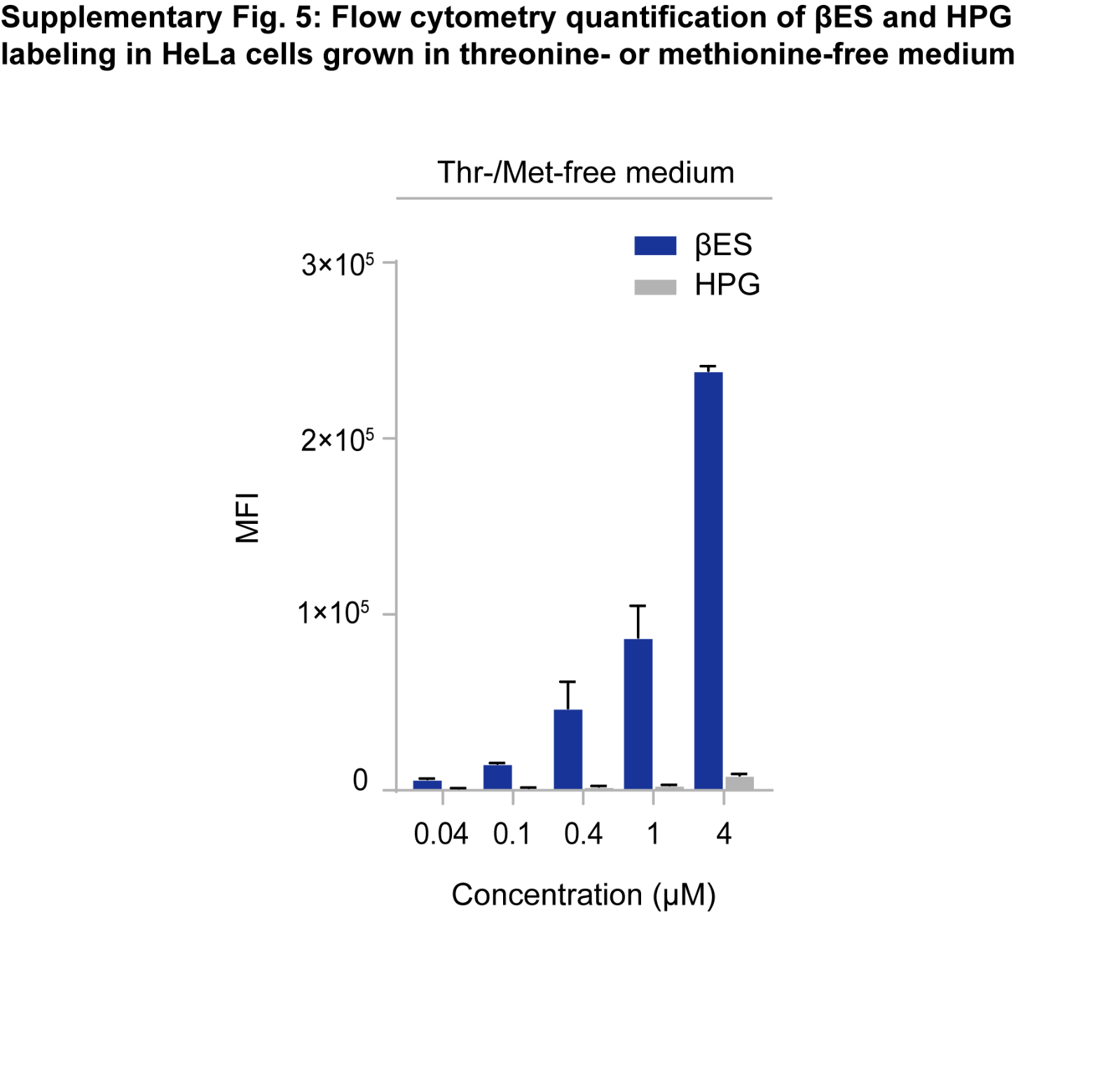

**Supplementary Fig. 5: Flow cytometry quantification of βES and HPG incorporation into HeLa proteome** **in threonine- or methionine-free medium.** HeLa cells were starved for 1 h in threonine-free or methionine-free medium and incubated for 1 h with indicated concentrations of βES in threonine-free medium or HPG in methionine-free medium, respectively. Incorporated analog was conjugated to Cy5-azide for quantification. MFI, mean fluorescence intensity; AU, arbitrary units; Thr, threonine; Met, methionine. Error bars represent s.d. Sample size is *n* = 3.

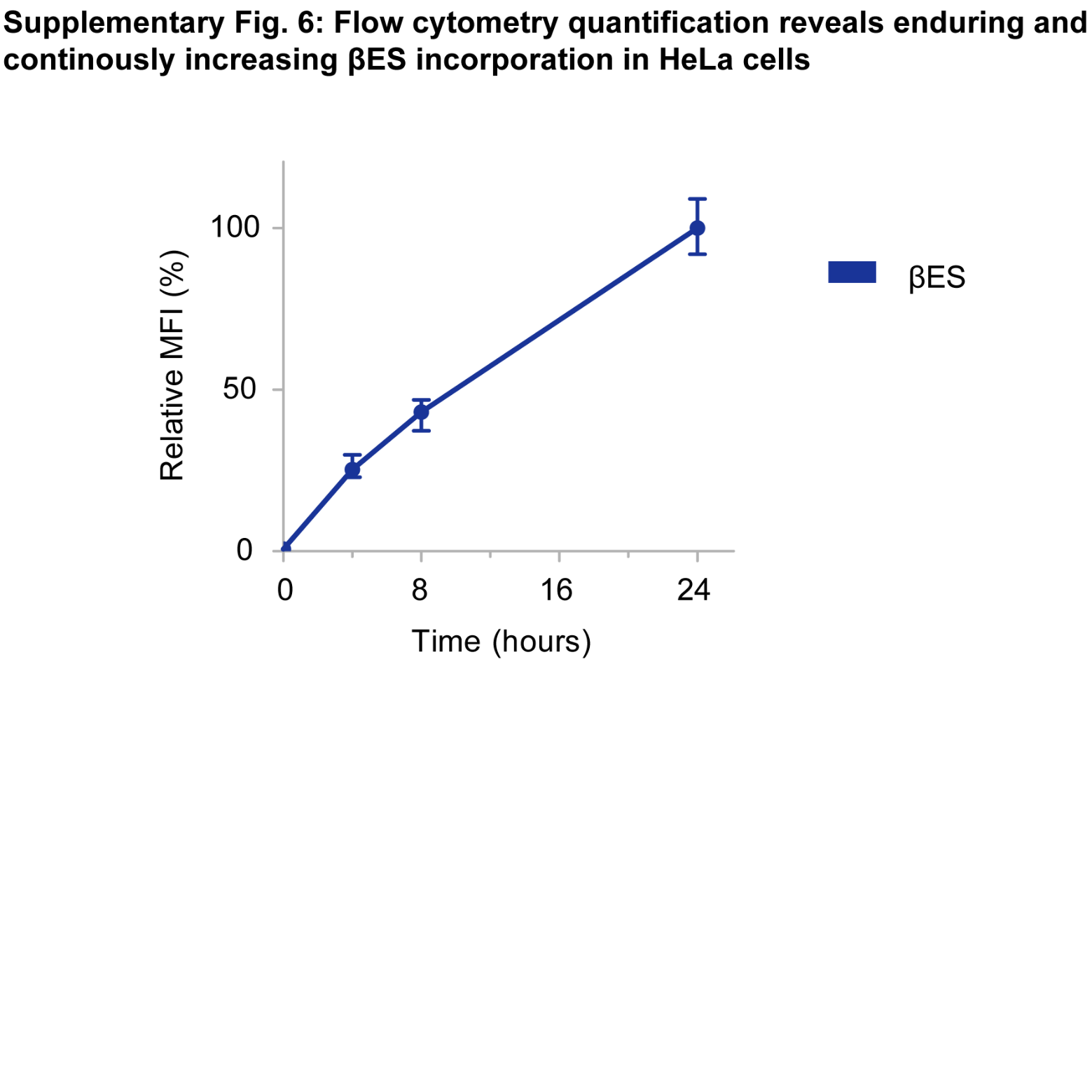

**Supplementary Fig. 6: Flow cytometry quantification of βES incorporation into HeLa proteome over time.** HeLa cells were incubated with 4 mM βES in complete medium for the indicated duration. Untreated HeLa cells were taken along as a control (0 h). Incorporated analog was conjugated to Cy5-azide for quantification. Signals are normalized to that at 24 h, which is set to 100%. MFI, Mean fluorescence intensity. Error bars represent s.d. Sample size is *n* = 3.

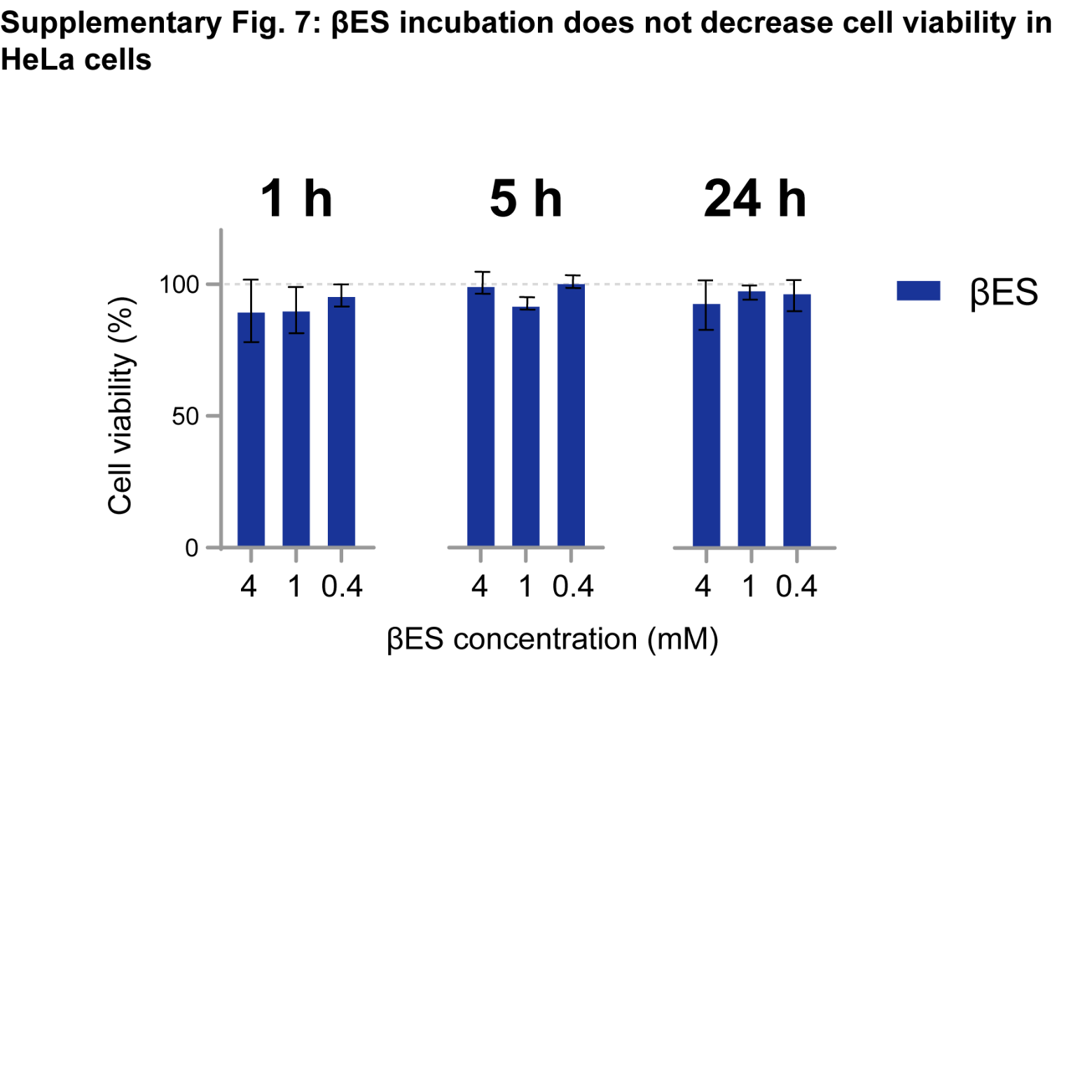

**Supplementary Fig. 7: βES incubation does not decrease cell viability in HeLa cells.** HeLa cells were incubated with indicated concentrations of βES in complete medium for the indicated duration. Cell viability was determined by exclusion of propidium iodide dye. Untreated HeLa cells were taken along as a control. Signals are normalized to that of control, which is set to 100%. Error bars represent s.d. Sample size is *n* = 4.

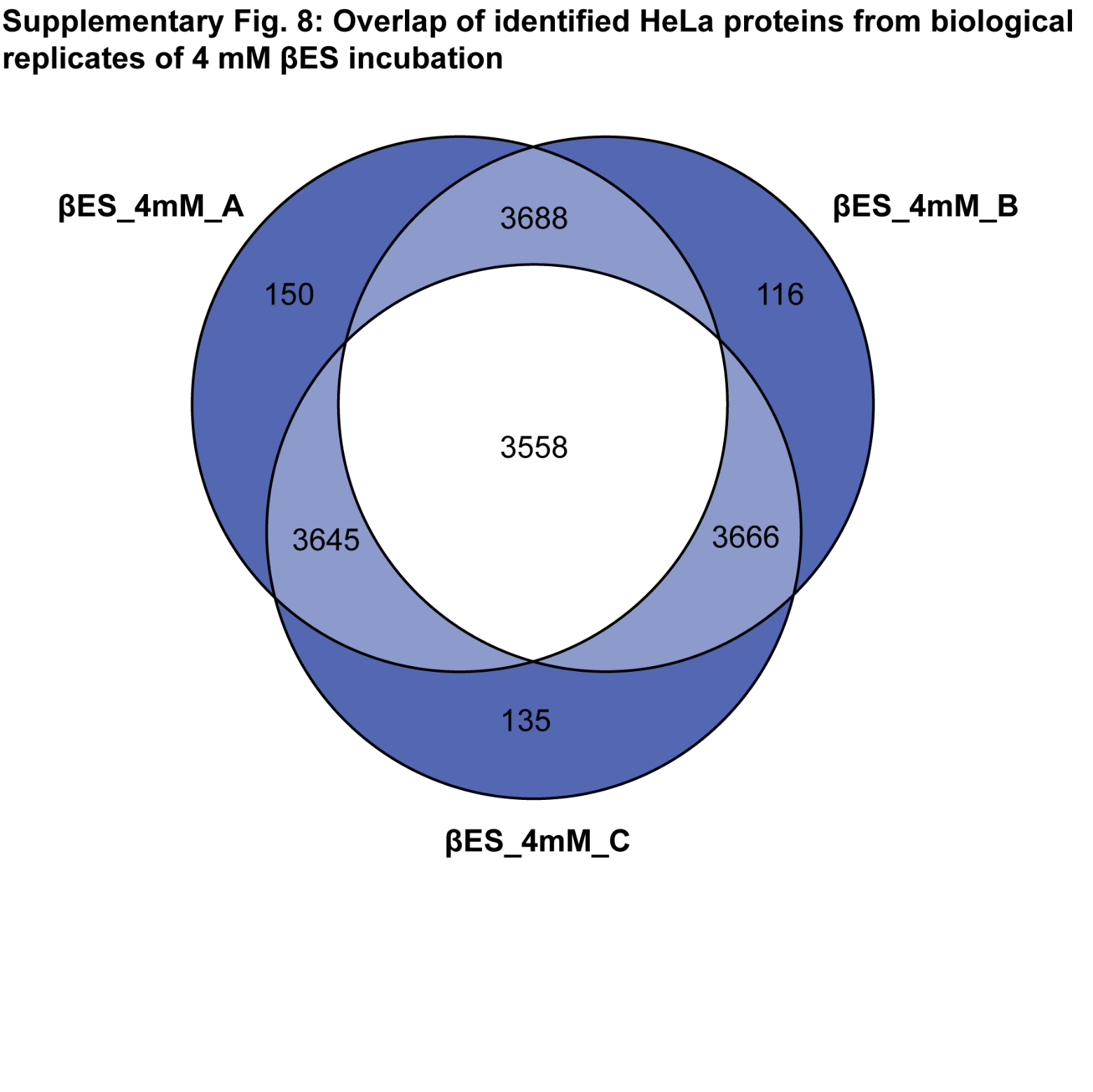

**Supplementary Fig. 8: Overlap between HeLa proteins identified in three replicate THRONCAT experiments.** HeLa cells were incubated for 5 h with 4 mM βES in complete medium. NSPs were enriched from cell lysate, digested and peptides subjected to LC-MS/MS analysis. Numbers in diagram indicate an amount of identified proteins. See Supplementary Data 1 for complete list of identified proteins.

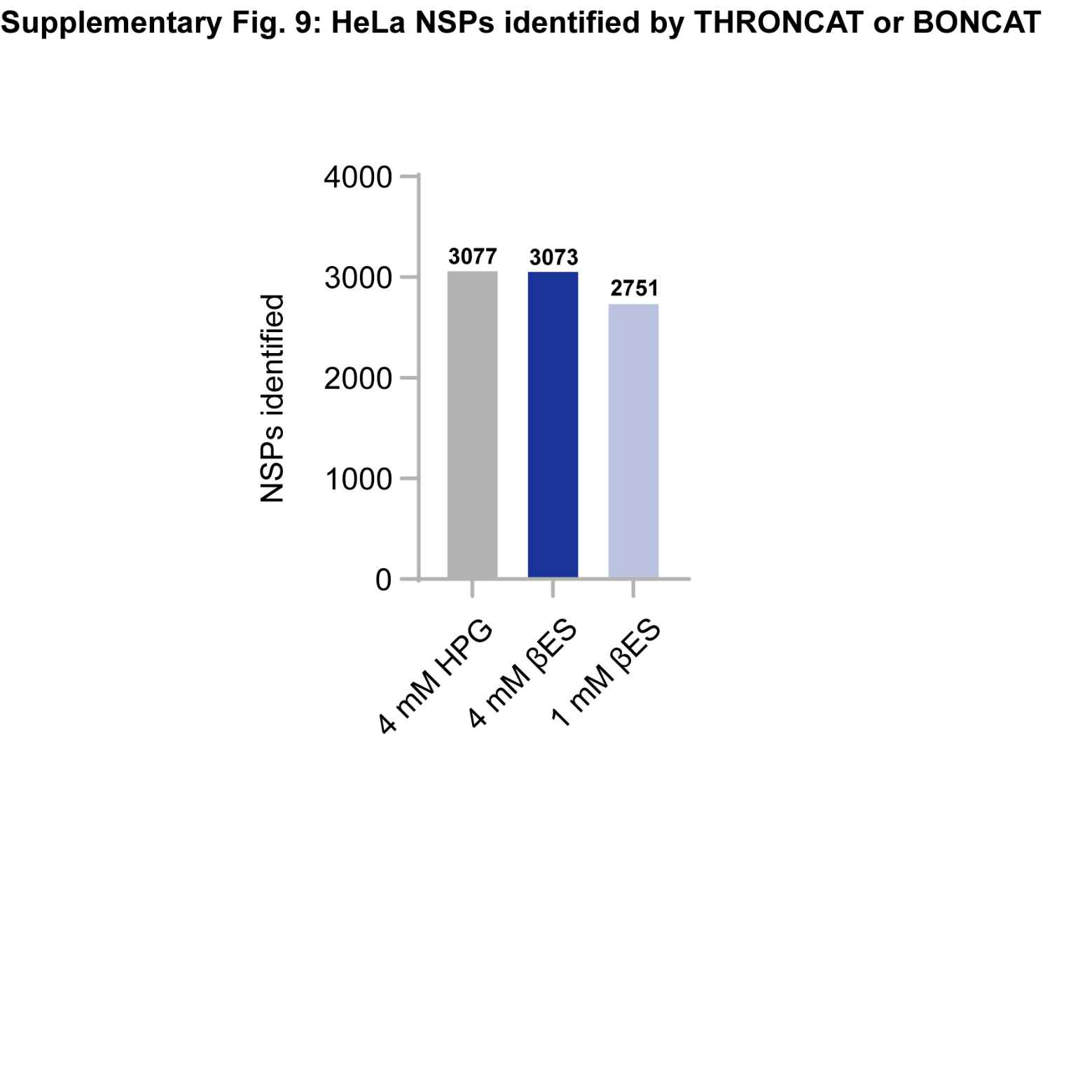

**Supplementary Fig. 9: NSPs confidently identified by THRONCAT and BONCAT.** HeLa cells were incubated for 5 h with 1 mM or 4 mM βES in complete medium (THRONCAT) or HeLa cells were starved for 1 h in methionine-free medium and incubated for 5 h in 4 mM HPG in methionine-free medium. Untreated HeLa cells were taken along as a control. For all conditions three biological replicates were performed. NSPs were enriched from cell lysate, digested and peptides subjected to LC-MS/MS analysis. Identified proteins were filtered for occurrence in all three replicates. To assign proteins as ‘confidently identified’ NSPs, proteins occurring in all three replicates of the control were subtracted.

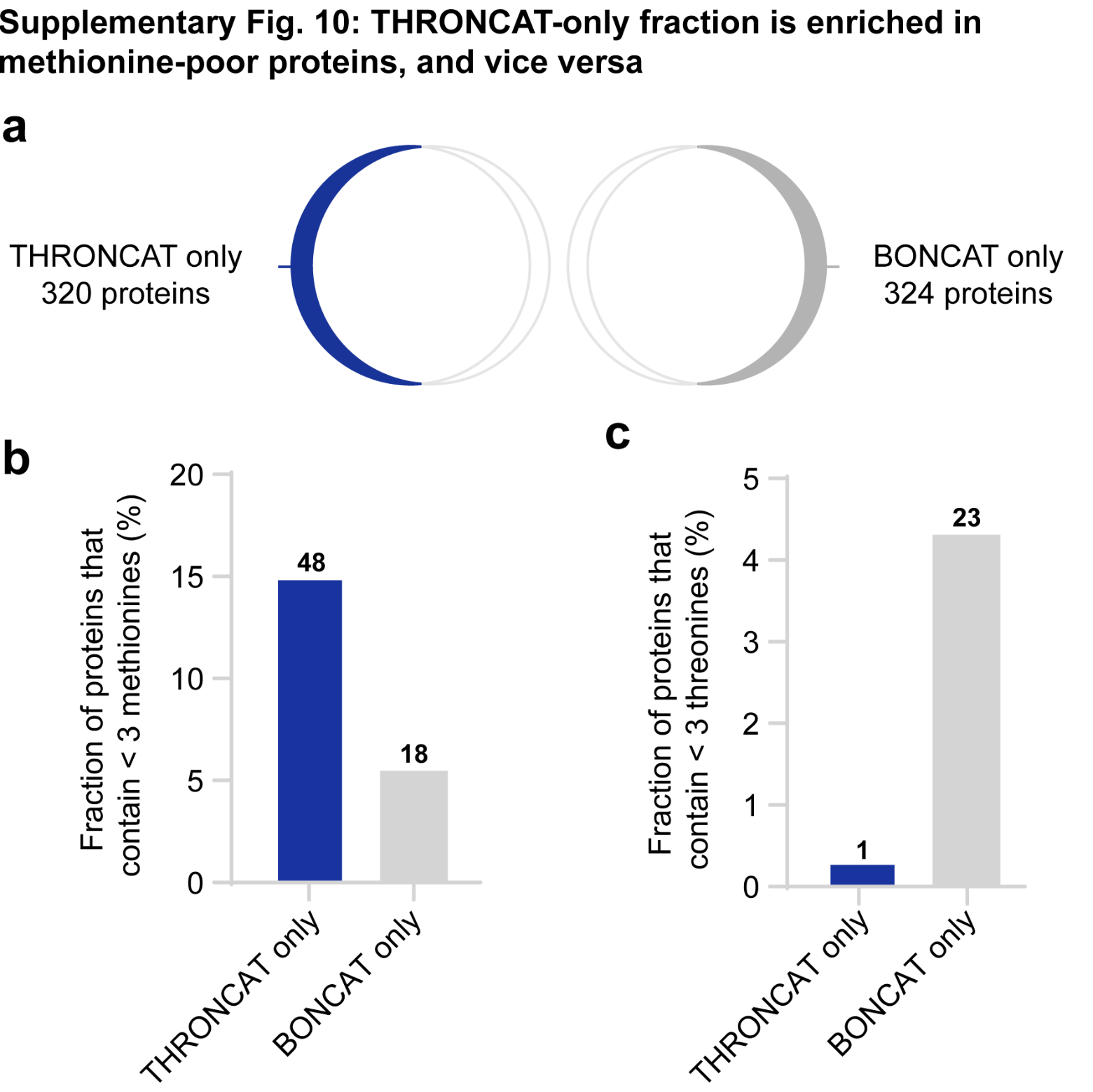

**Supplementary Fig. 10: Analysis of protein composition of THRONCAT-only and BONCAT-only fractions. a,** scheme showing THRONCAT-only and BONCAT-only fractions in venn diagram from **Fig. 4c**. **b,** Relative abundance of methionine-poor proteins in THRONCAT-only and BONCAT-only fractions. Methionine-poor proteins, containing less than 3 methionine residues, were enriched in the THRONCAT-only fraction. **c,** Relative abundance of threonine-poor proteins in THRONCAT-only and BONCAT-only fractions. Threonine-poor proteins, containing less than 3 threonine residues, were enriched in the BONCAT-only fraction. **b-c**, Protein composition was analyzed using python script AACALC (Supplementary Data 1). Numbers above bars represent the absolute amount of methionine-poor proteins identified in each fraction.

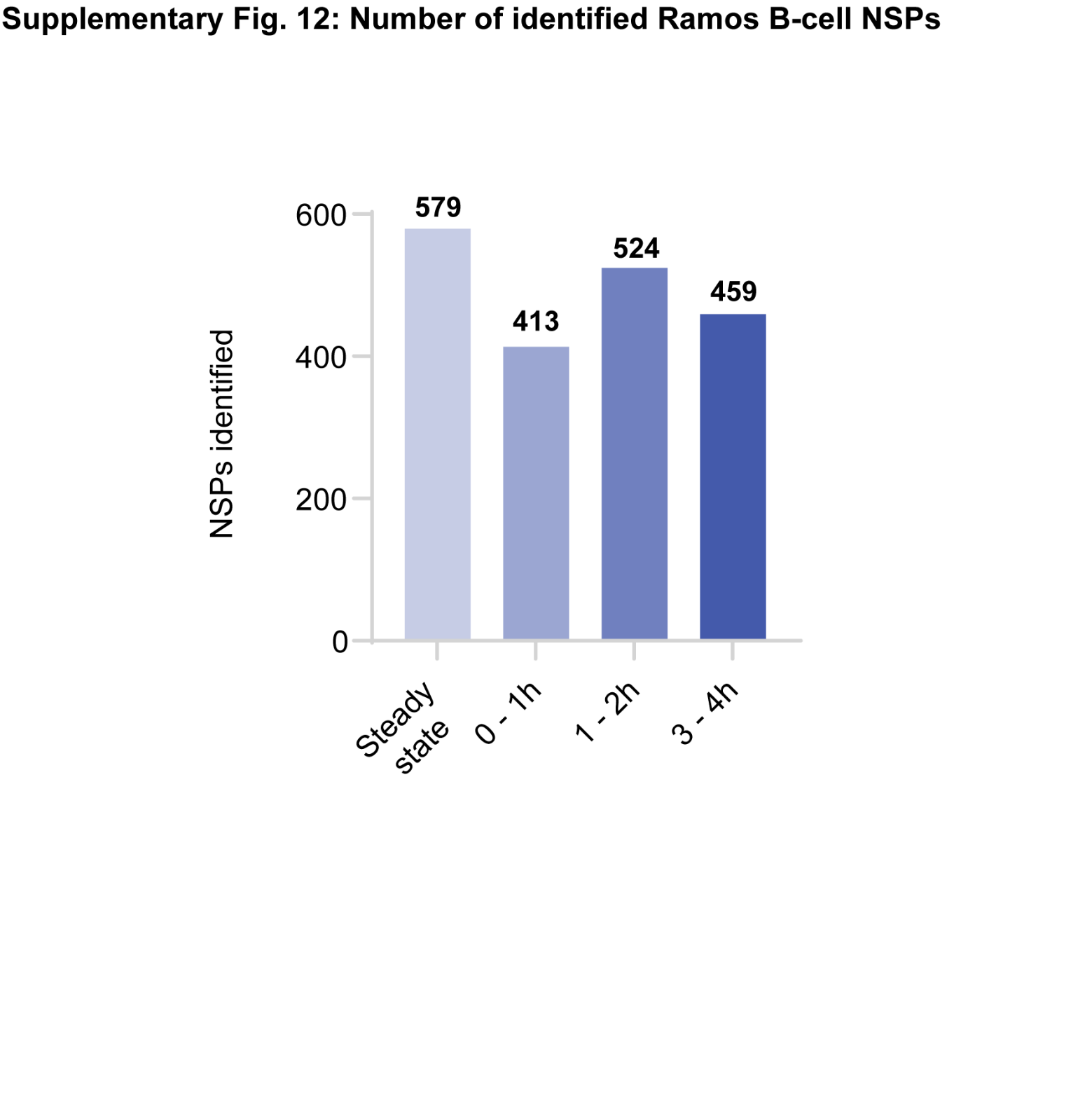

**Supplementary Fig. 11: NSPs confidently identified at different time points during Ramos cell activation.** Ramos B cells were pulse-labeled with 1 mM βES and d_8_-lysine for 1 h. NSPs were enriched, digested and subjected LC-MS/MS analysis. Only peptides containing d_8_-lysine were used for protein identification and NSPs were assigned as ‘confidently identified’ if they were present in all three biological replicates.

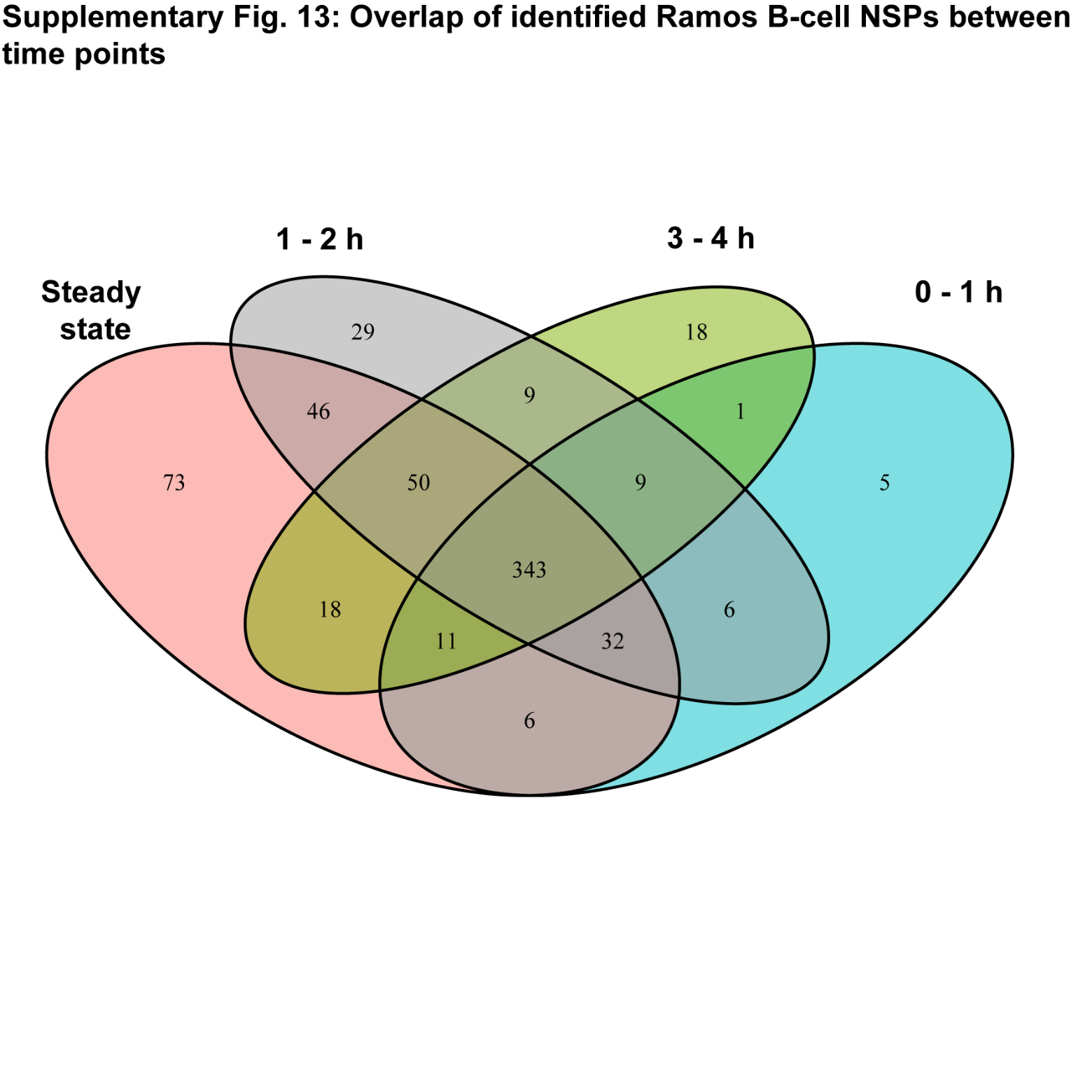

**Supplementary Fig. 12: Activation of Ramos B cells induces significant proteomic changes.** Venn diagram showing the overlap in number of NSPs identified at steady state and at different time points using during the activation of Ramos B cells by THRONCAT.
