## Supplementary Note 1 for "THRONCAT: Efficient metabolic labeling of newly synthesized proteins using a bioorthogonal threonine analog"

### **Supplementary Note 1: Detailed description of organic syntheses**

#### **General synthetic methods**

1H and 13C NMR spectra were recorded on a Bruker 400 MHz or 500 MHz spectrometer. Chemical shifts are reported in parts per million (ppm) relative to tetramethylsilane (TMS), or residual solvents as the internal standard. NMR data is presented as follows: chemical shift, multiplicity, coupling constant in hertz (Hz), integration. All NMR signals were assigned on the basis of ^1^H NMR, 13C APT NMR, COSY, HSQC and HMBC experiments. Mass spectra were recorded on an JEOL AccuTOF CS JMST100CS mass spectrometer. Automatic flash column chromatography was performed using a Biotage Isolera Spektra One. TLC-analysis was conducted on Silicagel F254 (Merck KGaA) with detection by UV-absorption (254nm) or staining with ninhydrin or KMnO_4_ solutions. DCM and THF were freshly distilled. All inert reactions were carried out under argon atmosphere using flame-dried flasks.

**(9H-fluoren-9-yl)methyl chlorocarbamate (5)**

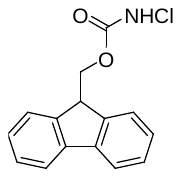

**5** was synthesized according to a previously described procedure.^1^ Briefly, 9-Fluorenylmethyl Carbamate (2.28 g, 9.55 mmol) was dissolved with heating in MeOH (170 mL). When the reaction mixture had cooled to 35 °C, trichloroisocyanuric acid (737 mg, 3.17 mmol) was added in one portion. The reaction mixture was stirred at room temperature for 18 hours, when another portion of trichloroisocyanuric acid (207 mg, 0.891 mmol) was added. After stirring for another 4.5 hours at room temperature, the reaction mixture was concentrated *in vacuo*. The resulting solid was suspended in hot toluene and then filtered while hot. Upon cooling, crystals appeared in the filtrate. The filtrate was heated and then allowed to slowly cool to room temperature, followed by cooling to 4 °C overnight. The resulting crystals were filtered and dried under high vacuum to yield **5** (2.52 g, 97%) as a white fluffy solid. **TLC** (EtOAc/*n*-heptane, 1/1, v/v): R_f_ = 0.68; **^1^H NMR** (400 MHz, CDCl_3_) δ 7.77 (dt, J = 7.6, 1.0 Hz, 2H), 7.61 (dq, J = 7.5, 1.0 Hz, 2H), 7.44 – 7.39 (m, 2H), 7.33 (td, J = 7.5, 1.2 Hz, 2H), 5.56 (s, 1H), 4.51 (d, J = 7.1 Hz, 2H), 4.26 (t, J = 6.9 Hz, 1H); **^13^C NMR** (101 MHz, CDCl_3_) δ 156.70, 143.22, 141.35, 127.97, 127.21, 125.05, 120.11, 70.44, 46.85; **HRMS** (m/z): [M + Na]^+^ calculated for C_15_H_12_ClNO_2_, 296.0454; found, 296.0455.

**Methyl (E)-5-(trimethylsilyl)pent-2-en-4-ynoate (2)**

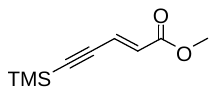

3-(Trimethylsilyl)-2-propynal (1.20 mL, 8.15 mmol) was dissolved in dry THF (35 mL) under inert atmosphere and cooled to 0 °C. Methyl (triphenylphosphoranylidene)acetate (3.28 g, 9.82 mmol) was added in one portion and the resulting mixture was stirred at 0 °C for 1.5 hours. The reaction mixture was concentrated *in vacuo* and silica column chromatography (0 – 10% Et_2_O in *n*-pentane) afforded the title compound **2** (1.41 g, 95%) as a clear liquid. **TLC** (EtOAc/*n*-heptane, 1/4, v/v): R_f_ = 0.67; **^1^H NMR** (400 MHz, CDCl_3_) δ 6.73 (d, J = 15.9 Hz, 1H), 6.24 (d, J = 16.0 Hz, 1H), 3.74 (s, 3H), 0.20 (s, 9H); **^13^C NMR** (101 MHz, CDCl_3_) δ 167.19, 133.40, 128.58, 105.49, 101.65, 52.71, -0.003; **HRMS** (m/z): [M + H]^+^ calculated for C_9_H_14_O_2_Si, 183.0841; found, 183.0835.

**Methyl (2S,3R)-2-((((9H-fluoren-9-yl)methoxy)carbonyl)amino)-3-hydroxy-5-(trimethylsilyl)pent-4-ynoate (3)**

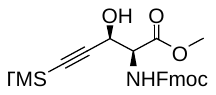

#### To a stirring suspension of **5** (3.21 g, 11.7 mmol) in *n*-PrOH (12 mL) cooled to 0 °C was added NaOH (359 mg, 8.98 mmol) in H_2_O (15 mL). To this mixture was added a solution of (DHQD)_2_AQN (601 mg, 0.704 mmol) in *n*-PrOH (18 mL), followed by a solution of **2** (1.09 g, 5.98 mmol) in *n*-PrOH (10 mL). At this point, the reaction mixture was a homogenous, bright yellow solution. A solution of K_2_OsO_4_ · 2 H_2_O (176 mg, 0.479 mmol) in H_2_O (5 mL, a few drops of the NaOH solution were added to the solution of K_2_OsO_4_ · 2 H_2_O) was added to the reaction mixture, upon which the color of the reaction mixture turned to deep green. The reaction mixture was stirred for 4 hours at °C when the color of the reaction mixture had turned from deep green to yellow again and TLC indicated completion. The reaction mixture was diluted with EtOAc (30 mL) and then washed with sat. aq. Na_2_S_2_O_3_ (50 mL), sat. aq. NaHCO_3_ (50 mL) and brine (50 mL). The organic layer was dried over MgSO_4_, filtered and concentrated *in vacuo*. Purification over silica column chromatography (10 – 25% EtOAc in *n­*-heptane), followed by recrystallization from *n*-heptane yielded title compound **3** (803 mg, 31%, 86% *ee*) as a white crystalline solid. **TLC** (EtOAc/n-heptane, 1/1, v/v): 0.55; **Specific rotation** [α]­_D_^20^ 21.9 (*c* 1.9, CH_2_Cl_2_); **^1^H NMR** (400 MHz, CDCl_3_) δ 7.77 (m, J = 7.6, 1.0 Hz, 2H), 7.62 (m, J = 7.6 Hz, 2H), 7.41 (m, J = 7.5, 1.0 Hz, 2H), 7.36 – 7.29 (m, 2H), 5.63 (d, J = 9.0 Hz, 1H), 4.81 (dd, J = 6.9, 3.4 Hz, 1H), 4.64 (dd, J = 9.1, 3.4 Hz, 1H), 4.41 (d, J = 7.3 Hz, 2H), 4.26 (t, J = 7.1 Hz, 1H), 3.81 (s, 3H), 2.74 (d, J = 6.8 Hz, 1H), 0.15 (s, 9H); **^13^C NMR** (101 MHz, CDCl_3_) δ 170.22, 144.07, 141.64, 128.10, 127.44, 125.46, 120.35, 67.82, 64.02, 58.82, 53.22, 47.43, 0.00; **HRMS** (m/z): [M + Na]^+^ calculated for C_24_H_27_NO_5_Si, 460.1556; found, 460.1554.

**Methyl (2S,3R)-2-((((9H-fluoren-9-yl)methoxy)carbonyl)amino)-3-hydroxypent-4-ynoate (4)**

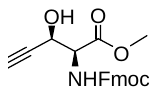

To a stirring solution of **3** (806 mg, 1.84 mmol) in DCM (15 mL) cooled to 0 °C was added TBAF (1M solution in THF; 3.68 mL, 3.68 mmol) dropwise. The reaction mixture was stirred for 10 minutes at 0 °C, when TLC indicated full conversion of **3**. The reaction mixture was diluted with EtOAc (10 mL) and washed with sat. aq. NH_4_Cl (2 x 25 mL) and brine (25 mL), dried over MgSO_4_, filtered and concentrated *in vacuo*. Purification over silica column chromatography (10 – 30 % EtOAc in *n*-heptane yielded title compound **4** (549 mg, 82 %) as a white crystalline solid. **TLC** (EtOAc/n-heptane, 1/1, v/v): R_f_ = 0.35; **Specific rotation** [α]­_D_^20^ 4.5 (*c* 2.2, CH_2_Cl_2_) **^1^H NMR** (400 MHz, CDCl_3_) δ 7.77 (dt, J = 7.6, 0.9 Hz, 2H), 7.62 (d, J = 7.5 Hz, 2H), 7.44 – 7.37 (m, 2H), 7.36 – 7.28 (m, 2H), 5.70 (d, J = 9.2 Hz, 1H), 4.86 (s, 1H), 4.66 (dd, J = 9.5, 3.1 Hz, 1H), 4.50 – 4.32 (m, 2H), 4.26 (t, J = 7.1 Hz, 1H), 4.12 (q, J = 7.2 Hz, 0H), 3.81 (s, 3H), 3.04 (s, 1H), 2.49 (d, J = 2.2 Hz, 1H); **^13^C NMR** (101 MHz, CDCl_3_) δ 156.24, 143.70, 141.31, 127.76, 127.09, 125.12, 120.02, 80.52, 75.06, 63.05, 58.31, 53.01, 47.09, 30.95, 22.71, 14.13; **HRMS** (m/z): [M + Na]^+^ calculated for C_21_H_19_NO_5_, 388.1161; found, 388.1156.

**(2S,3R)-2-amino-3-hydroxypent-4-ynoic acid, β­-ethynylserine (1)**

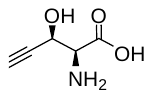

To a stirring solution of **3** (526 mg, 1.44 mmol) in MeCN (10 mL) cooled to 0 °C was added a solution of LiOH (172 mg, 7.20 mmol) in H_2_O (10 mL) dropwise. The reaction mixture was stirred at 0 °C for 6 hours, when TLC indicated complete deprotection of the methyl ester and Fmoc-group. The mixture was filtered through a C18-funtionalized silica plug (Screening Devices) Dowex50wx8 (Sigma, 10 mL) was washed with MeOH (50 mL) and H_2_O (50 mL), added to the reaction mixture and stirred at room temperature overnight. The mixture was filtered and the Dowex beads washed with 20 mL of a 3 M aq. NH_4_OH solution. The combined filtrate was concentrated *in vacuo* to yield title compound **1** (223 mg, 93%) as a pale yellow sticky solid. **TLC** (EtOAc/*n*-heptane, 1/1, v/v): **Specific rotation** [α]­_D_^20^ – 52.9 (*c* 1.2, H_2_O); R_f_ = **^1^H NMR** (500 MHz, D_2_O) δ 5.12 (ddd, J = 3.0, 2.2, 0.8 Hz, 1H), 4.29 (dd, J = 3.1, 0.8 Hz, 1H), 3.12 (dd, J = 2.4, 1.0 Hz, 1H). **^13^C NMR** (126 MHz, D_2_O) δ 168.84, 78.67, 77.55, 59.47, 57.58. **HRMS** (m/z): [M + Na]^+^ calculated for C_5_H_7_NO_3_, 152.0324; found, 152.0346.

##### **Mosher’s ester analysis for determination of absolute configuration of** **3**.^2^

To a stirred solution of **3** (9.1 mg, 20.8 μmol) in dry DCM (1 mL) was added, in this order, a catalytic amount of 4-dimethylaminopyridine (0.51 mg, 4.2 μmol), dry triethylamine (8.7 μL, 62 μmol) and *S*-(+)-MTPA-Cl (15.7 mg, 62 μmol). The reaction mixture was stirred at room temperature overnight, when TLC indicated full conversion of **3**. The reaction mixture was quenched by addition of sat. aq. NH_4_Cl (5 mL) and diluted with DCM (5 mL). The organic layer was washed with brine (5 mL), dried over MgSO_4_, filtered and dried *in vacuo*. Purification over silica column chromatography (0 – 20 % EtOAc in *n*-heptane) yielded the *R*-MTPA ester of **3** (7.4 mg, 55%) as a clear oil. **^1^H NMR** (500 MHz, CDCl_3_) δ 7.77 (ddt, J = 7.6, 3.2, 0.9 Hz, 2H), 7.59 (ddd, J = 11.8, 7.6, 3.1 Hz, 2H), 7.53 – 7.47 (m, 2H), 7.47 – 7.36 (m, 5H), 7.34 – 7.28 (m, 2H), 5.98 (d, J = 3.1 Hz, 1H), 5.50 (d, J = 9.7 Hz, 1H), 4.91 (dd, J = 9.7, 3.1 Hz, 1H), 4.37 (d, J = 7.5 Hz, 2H), 4.26 (dt, J = 14.7, 7.3 Hz, 1H), 3.74 (s, 3H), 3.54 – 3.50 (m, 3H), 0.10 (s, 9H). In an entirely analogues fashion, the *S*-MTPA ester of **3** was prepared from *R*-(-)-MPTA-Cl: **^1^H NMR** (500 MHz, CDCl_3_) δ 7.77 (dt, J = 7.6, 1.0 Hz, 2H), 7.62 – 7.55 (m, 2H), 7.55 – 7.48 (m, 2H), 7.48 – 7.37 (m, 5H), 7.30 (tdd, J = 7.5, 2.0, 1.2 Hz, 2H), 6.03 (d, J = 3.1 Hz, 1H), 5.45 (d, J = 9.7 Hz, 1H), 4.91 (dd, J = 9.7, 3.1 Hz, 1H), 4.39 – 4.34 (m, 2H), 4.26 (t, J = 7.5 Hz, 1H), 3.67 (s, 3H), 3.56 (s, 3H), 0.13 (s, 9H).

#### **NMR spectra**

400 MHz, CDCl_3_

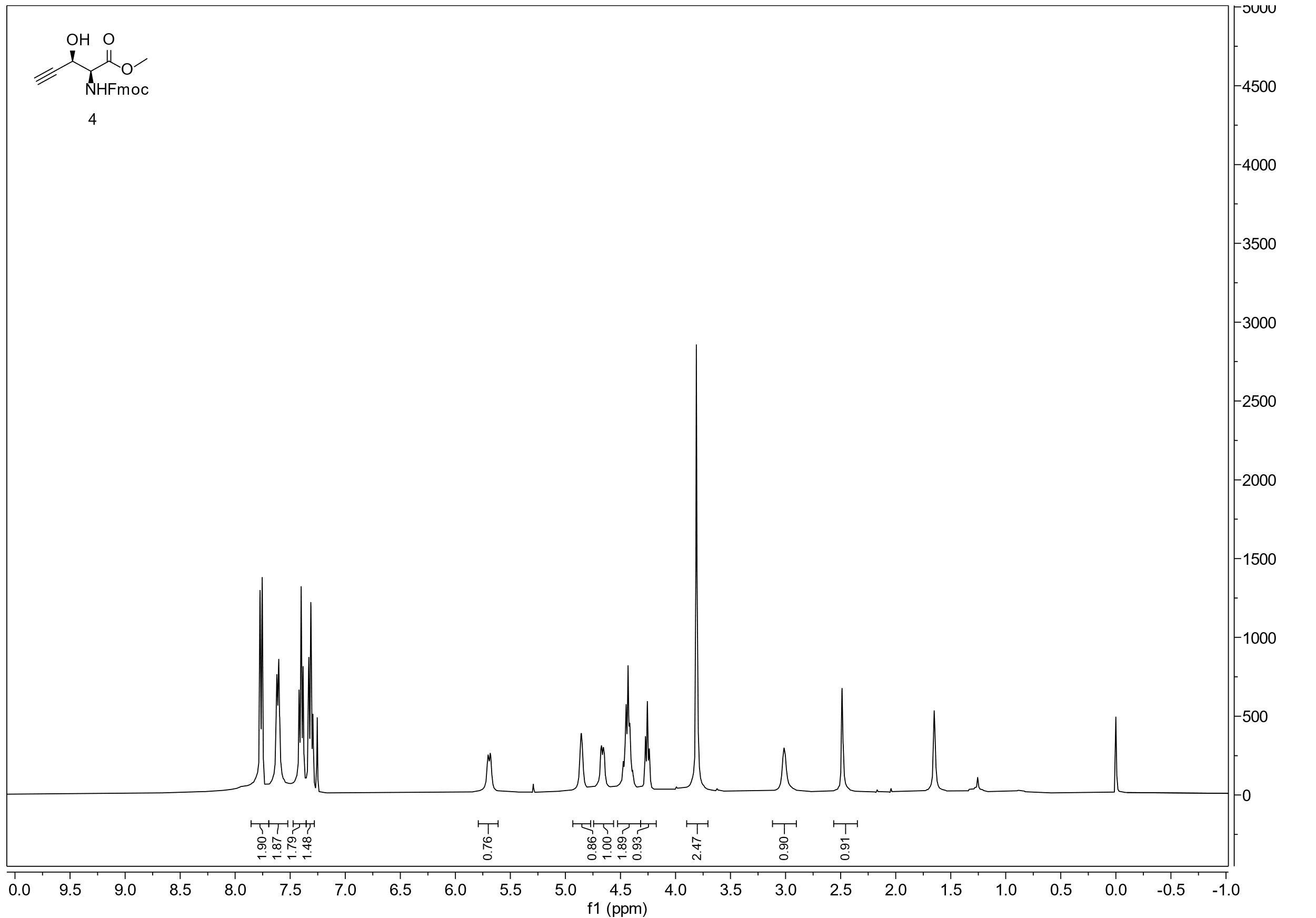

101 MHz, CDCl_3_

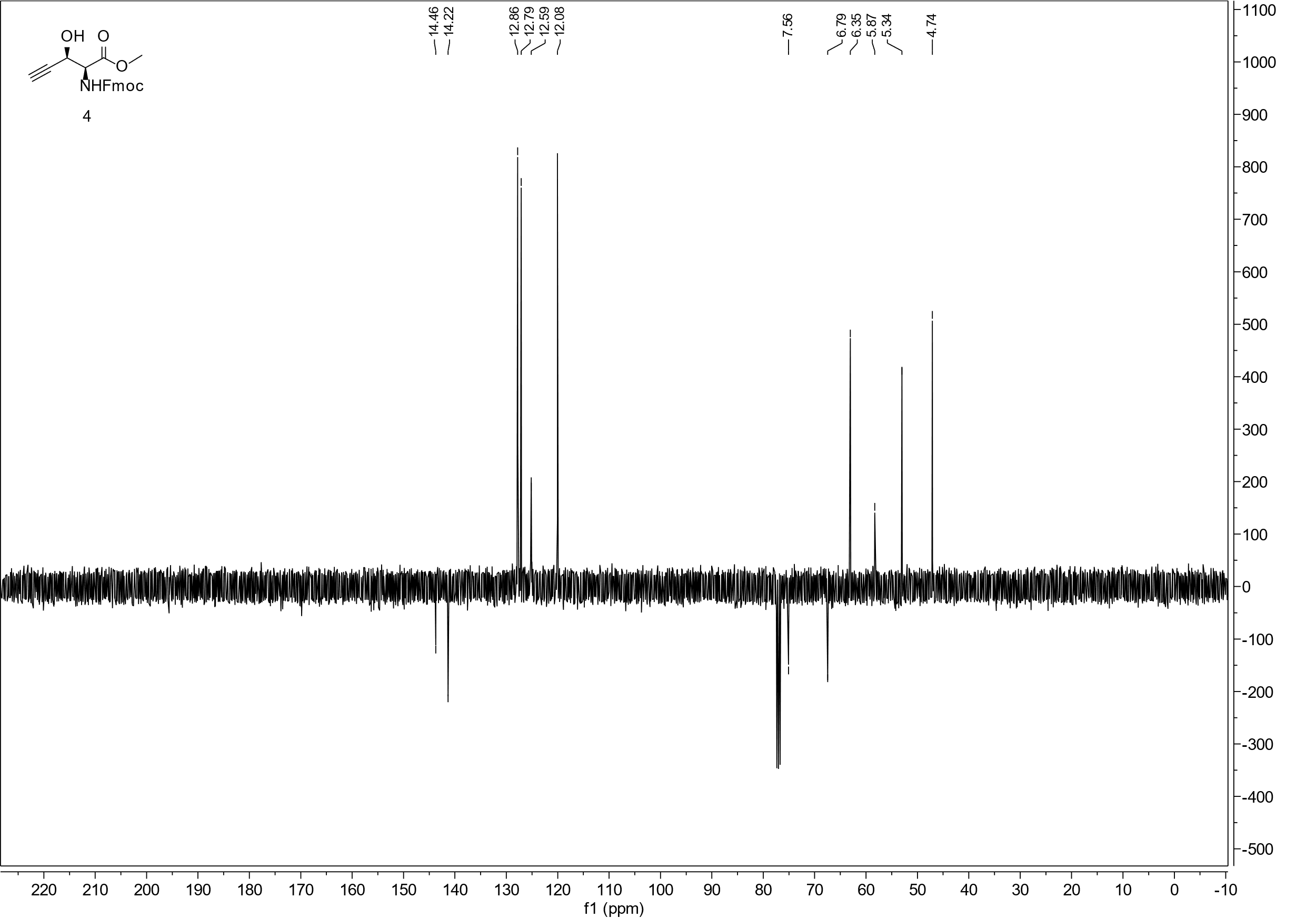

400 MHz, CDCl_3_

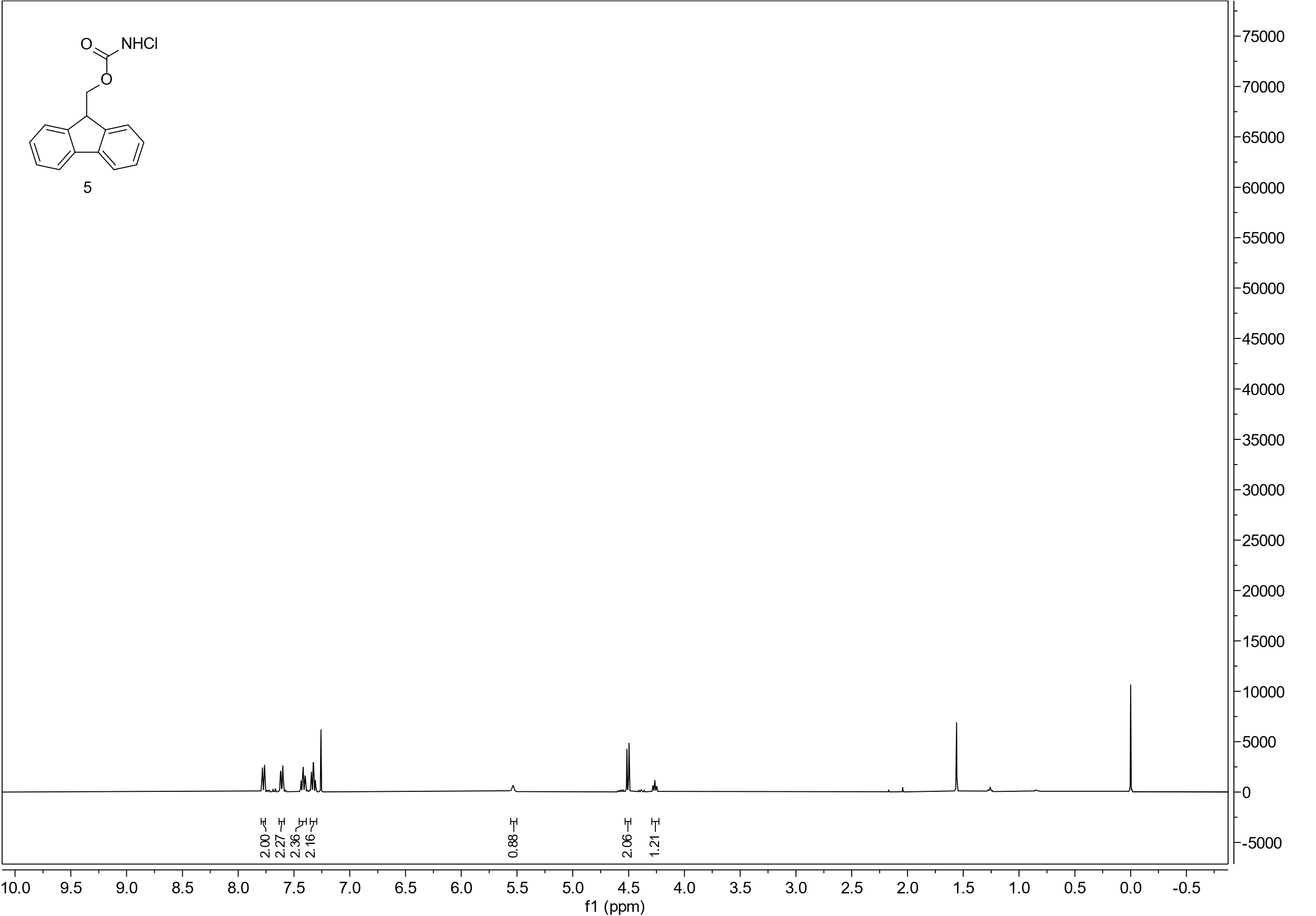

101 MHz, CDCl_3_

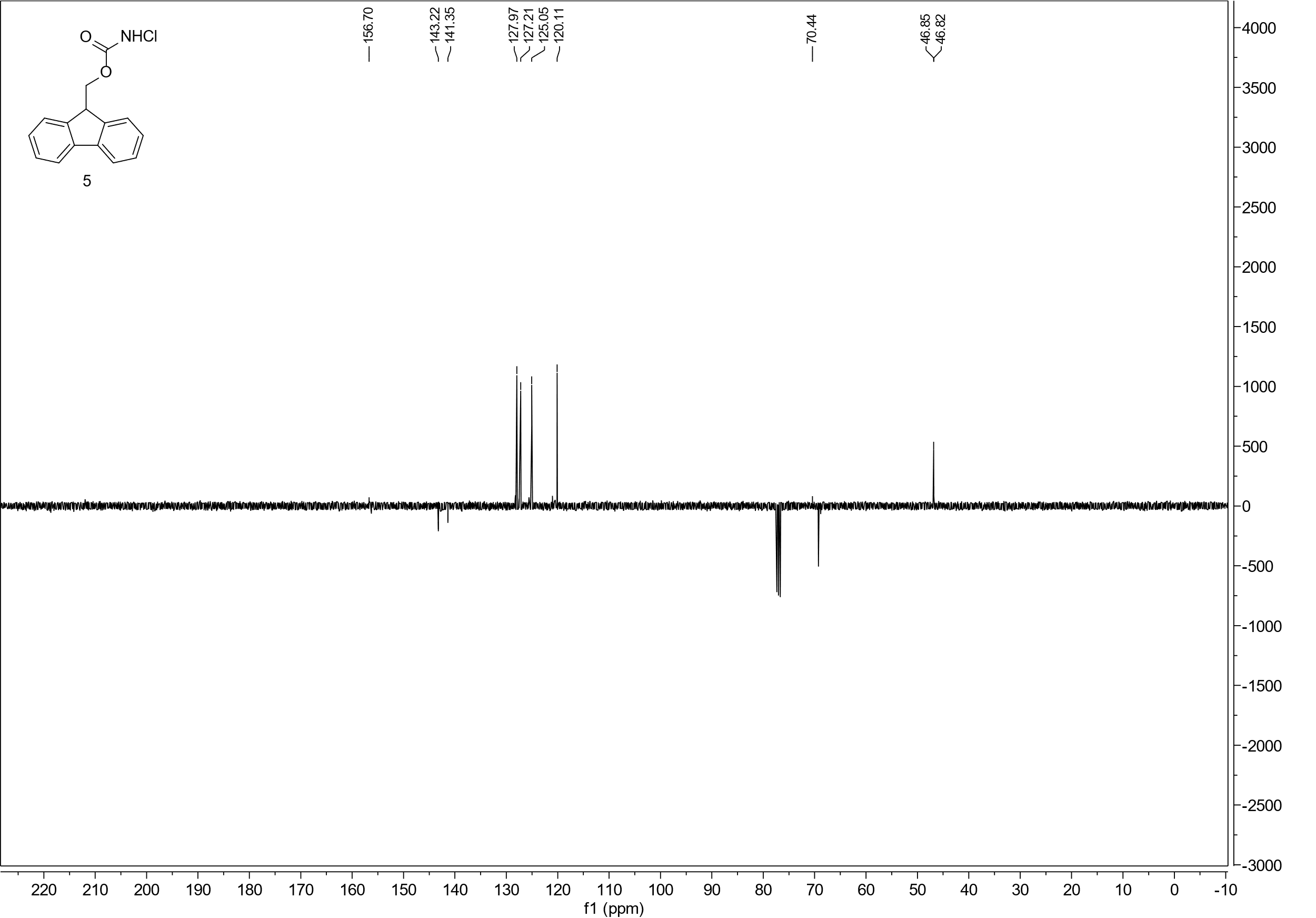

400 MHz, CDCl_3_

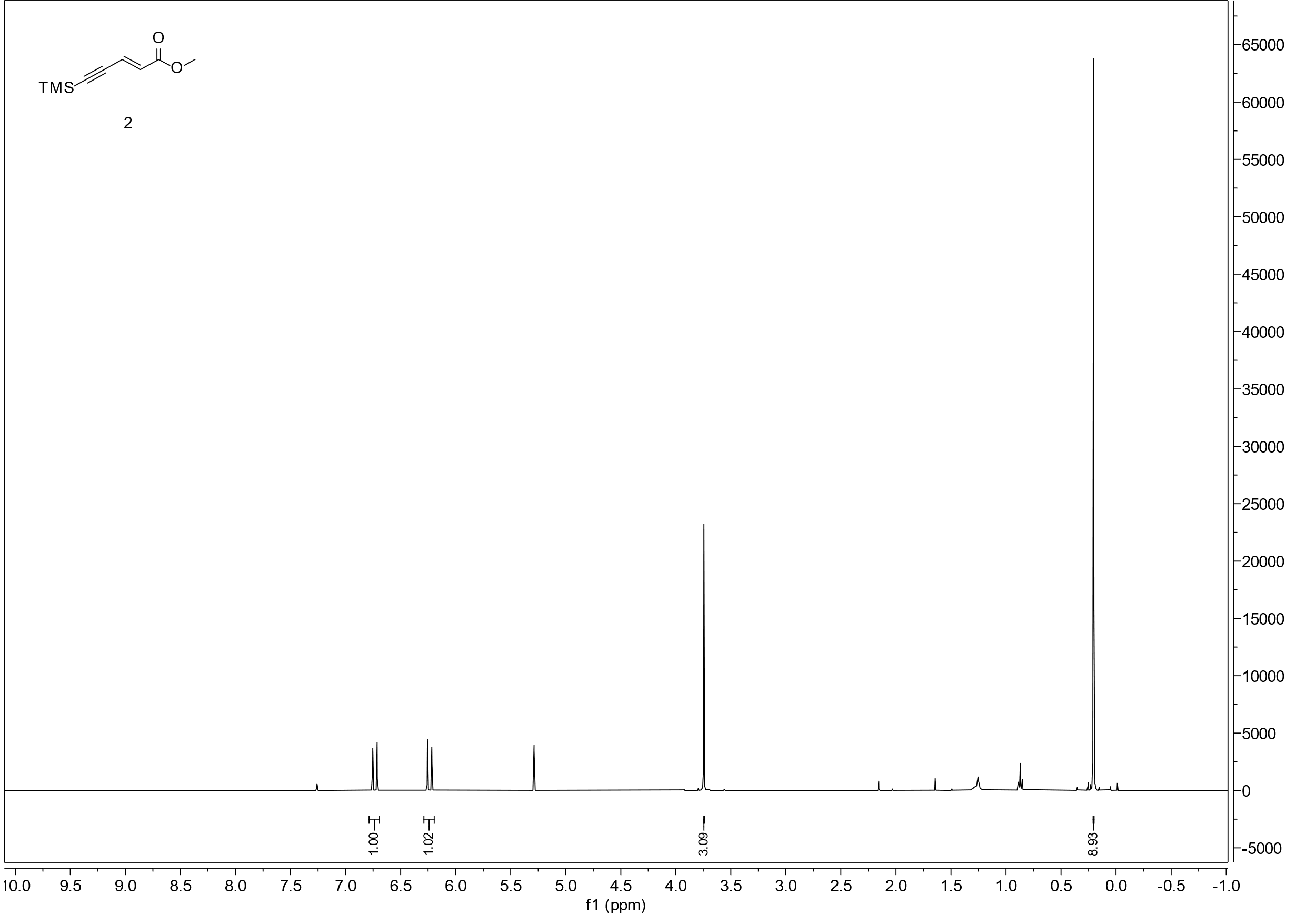

101 MHz, CDCl_3_

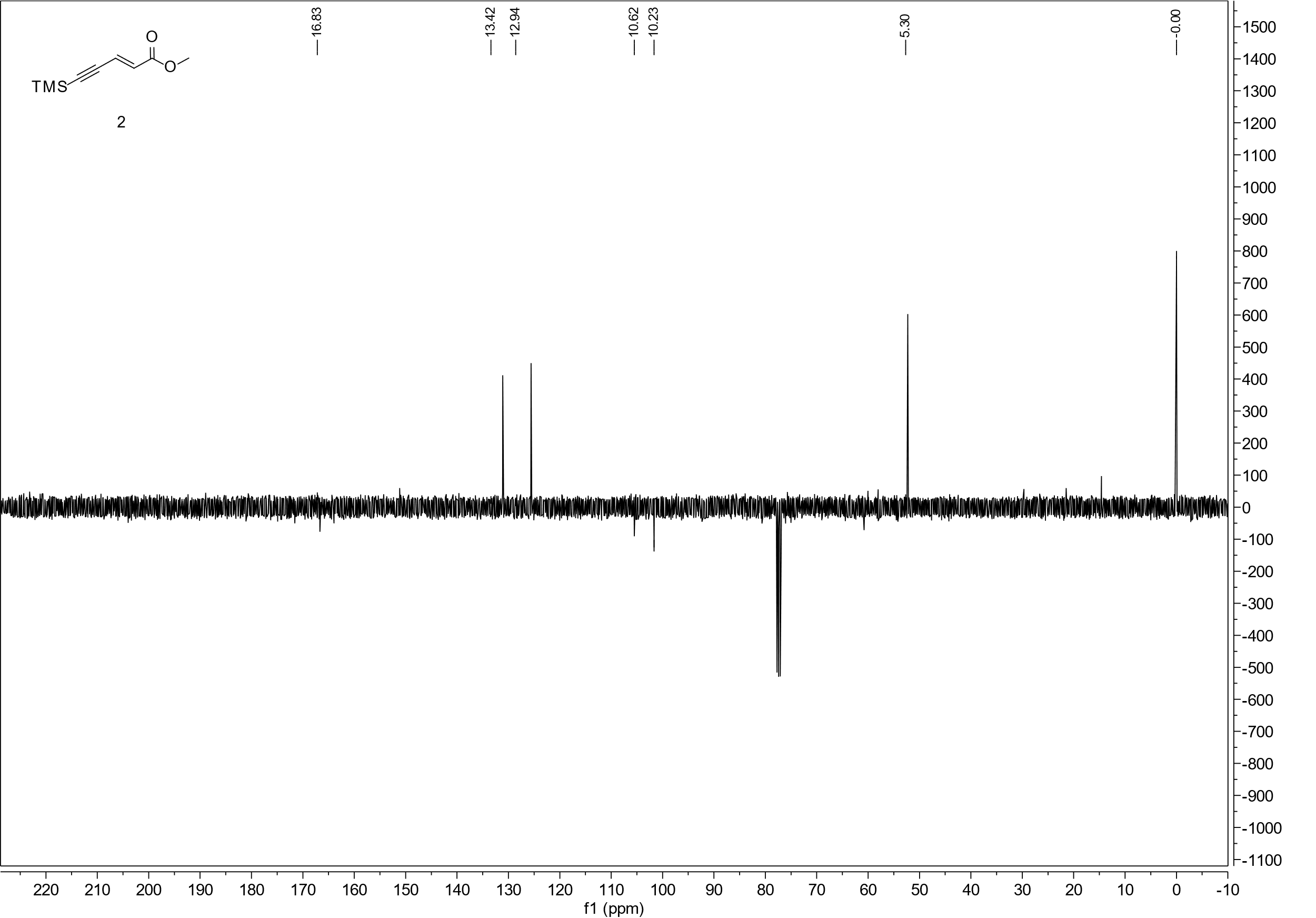

400 MHz, CDCl_3_

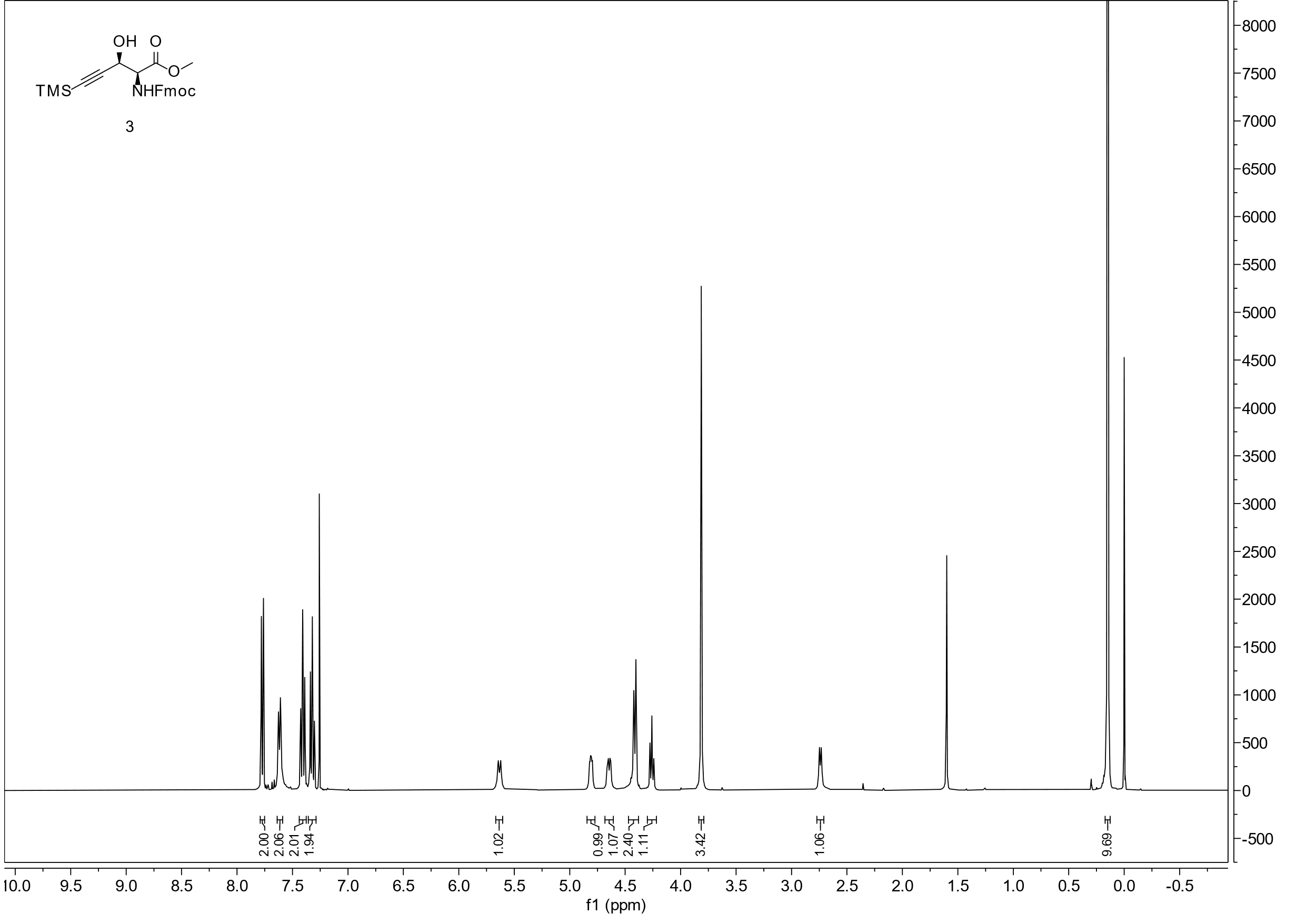

101 MHz, CDCl_3_

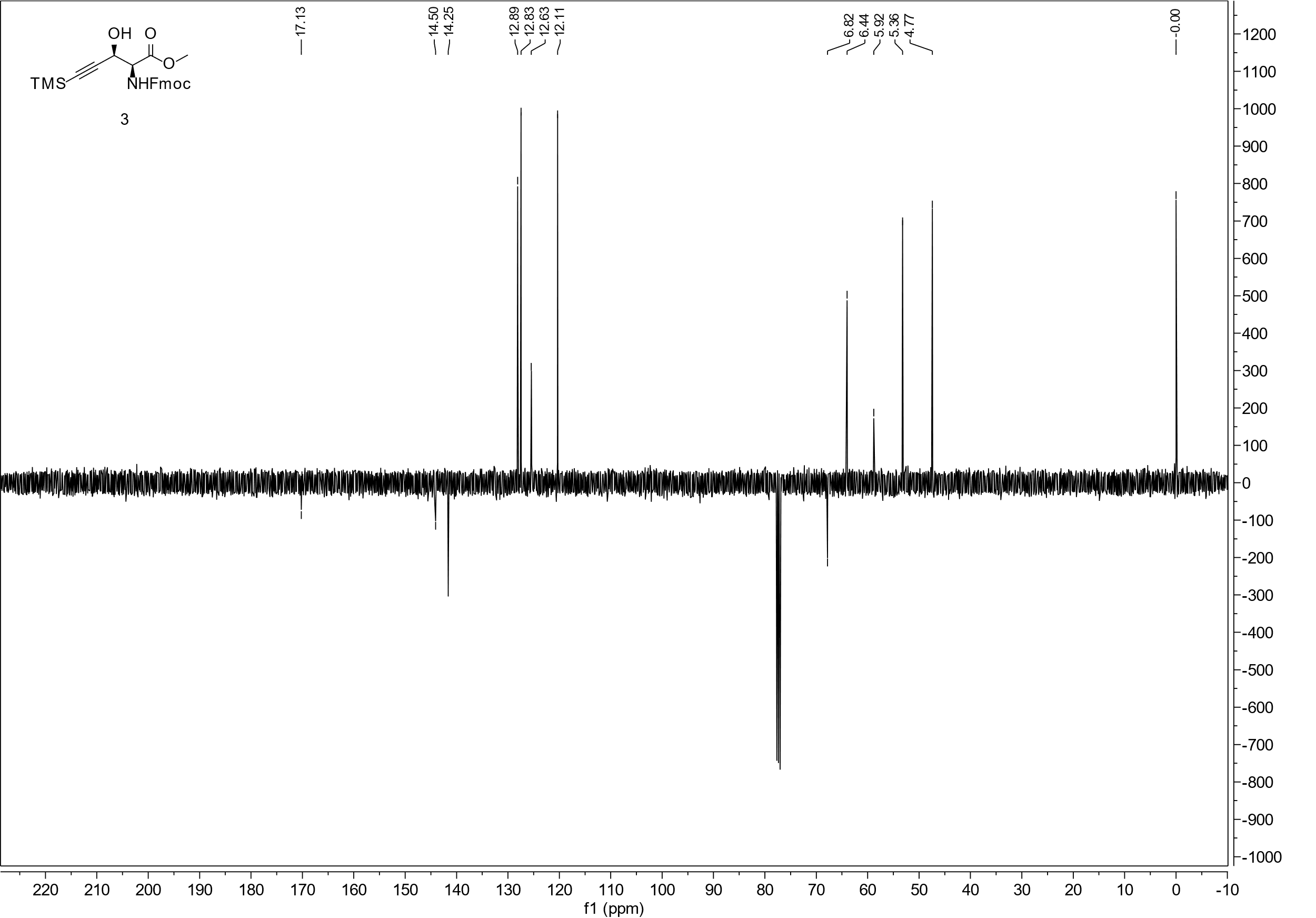

500 MHz, D_2_O

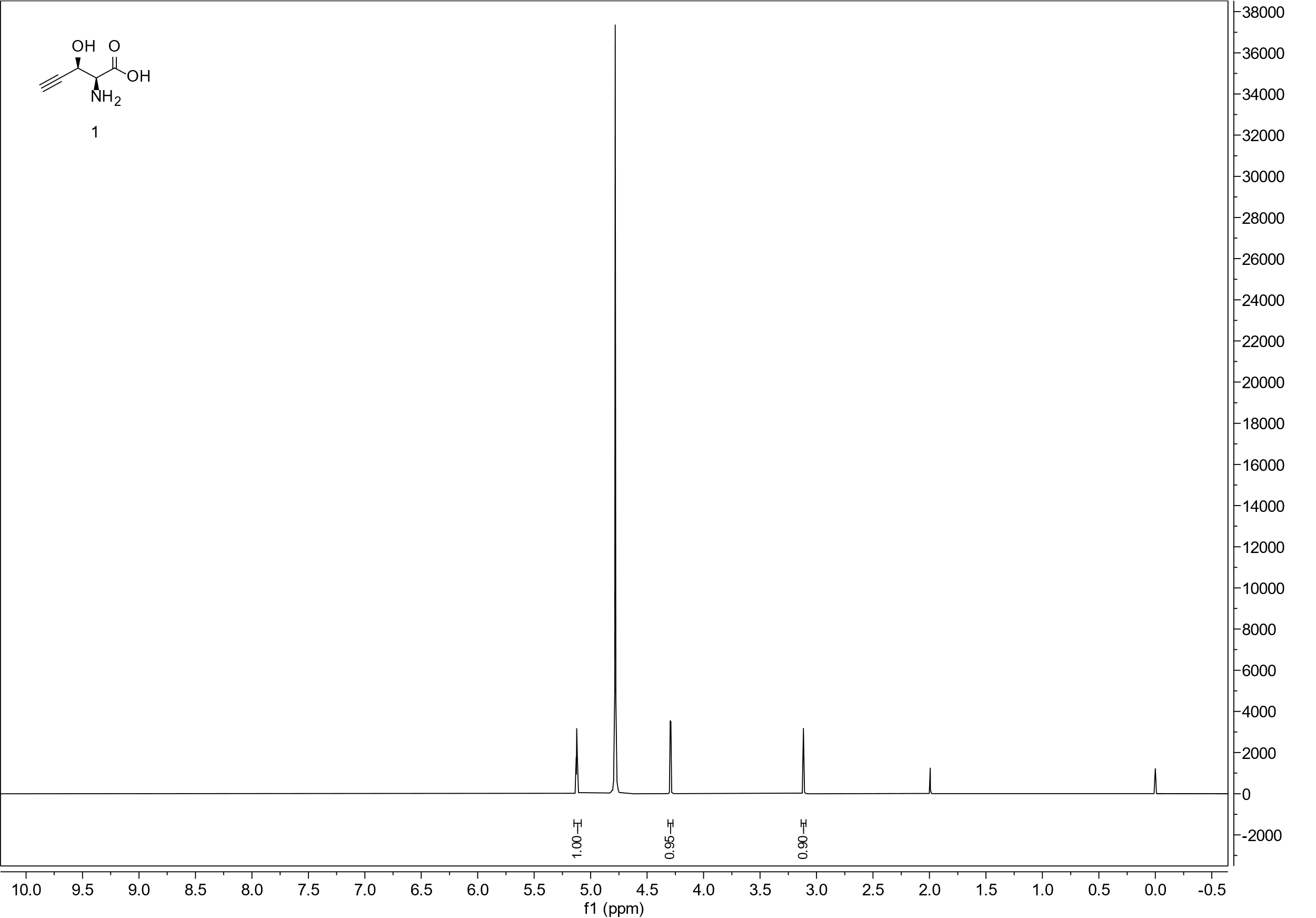

126 MHz, D_2_O

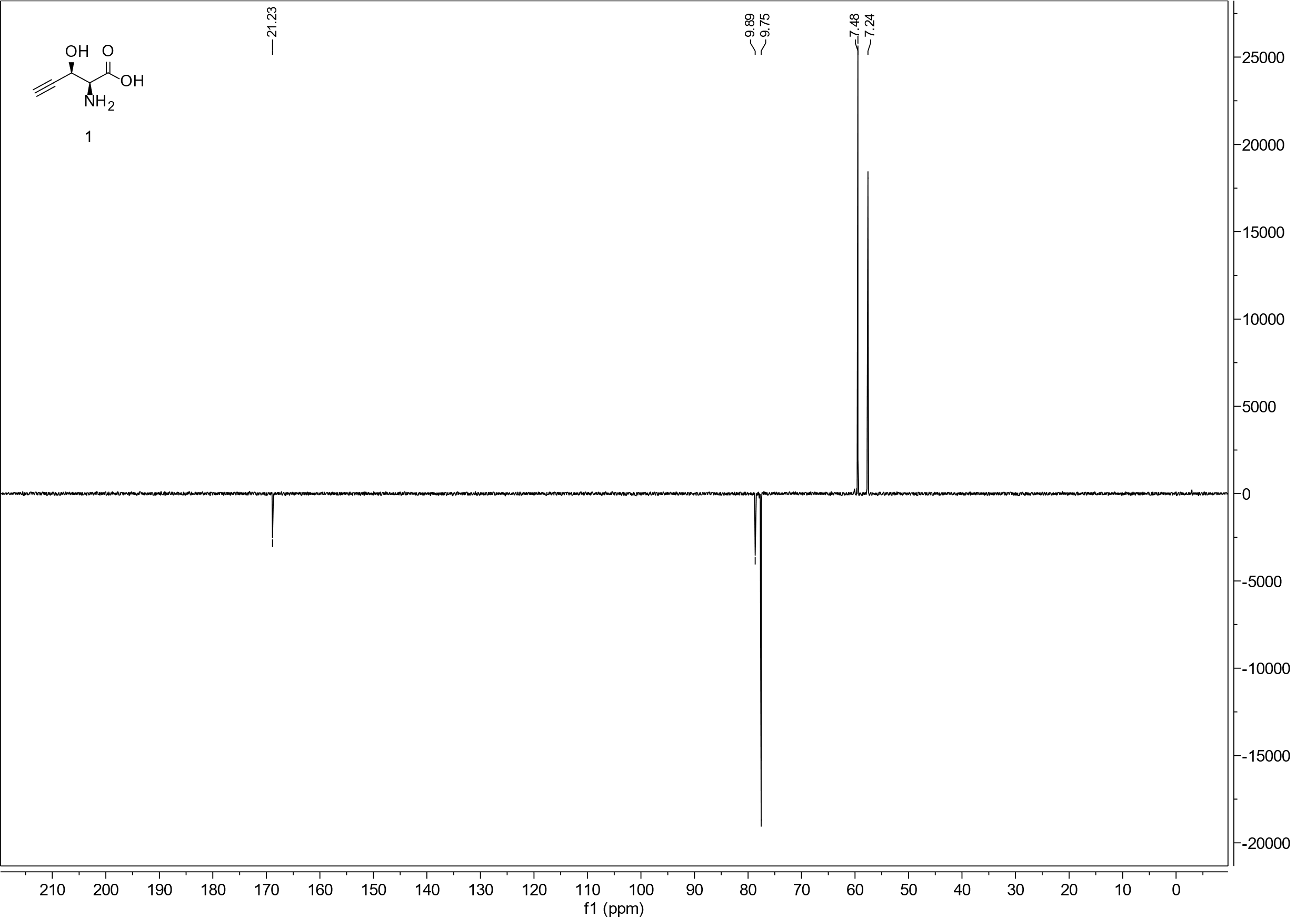

### Supplementary References

1.	Moreira, R. & Taylor, S. D. Asymmetric Synthesis of Fmoc-Protected β-Hydroxy and β-Methoxy Amino Acids via a Sharpless Aminohydroxylation Reaction Using FmocNHCl. *Org. Lett.* **20**, 7717–7720 (2018).

2.	Hoye, T. R., Jeffrey, C. S. & Shao, F. Mosher ester analysis for the determination of absolute configuration of stereogenic (chiral) carbinol carbons. *Nat. Protoc.* **2**, 2451–2458 (2007).
