## Supplementary Table 1 for "THRONCAT: Efficient metabolic labeling of newly synthesized proteins using a bioorthogonal threonine analog"

| Treatment | Time points | p-value | p-value summary |
| --- | --- | --- | --- |
| βES | 2h versus 4h | 0.012 | * |
|  | 2h versus 8h | 0.0012 | *** |
|  | 2h versus 16h | <0.0001 | *** |
|  | 2h versus 48h | <0.0001 | *** |
|  | 4h versus 8h | 0.37 | ns |
|  | 4h versus 16h | <0.0001 | *** |
|  | 4h versus 48h | <0.0001 | *** |
|  | 8h versus 16h | 0.0001 | *** |
|  | 8h versus 48h | <0.0001 | *** |
|  | 16h versus 48h | <0.0001 | *** |
| HPG | 2h versus 4h | >0.99 | ns |
|  | 2h versus 8h | 0.24 | ns |
|  | 2h versus 16h | 0.0004 | *** |
|  | 2h versus 48h | <0.0001 | *** |
|  | 4h versus 8h | >0.99 | ns |
|  | 4h versus 16h | 0.020 | * |
|  | 4h versus 48h | 0.0004 | *** |
|  | 8h versus 16h | 0.52 | ns |
|  | 8h versus 48h | 0.033 | * |
|  | 16h versus 48h | >0.99 | ns |

**Table S1:** Statistics to show that THRONCAT signal intensity increases with increasing labeling times. Brown-Forsythe ANOVA with Dunnett’s T3 multiple comparisons test was performed for βES time points. Kruskal-Wallis test with Dunn’s multiple comparisons test was performed for HPG time points. ns, not significant.
