## Supplementary Table 2 for "THRONCAT: Efficient metabolic labeling of newly synthesized proteins using a bioorthogonal threonine analog"

|  | **Threonine-free medium** | | **Methionine-free medium** | |
| --- | --- | --- | --- | --- |
|  | Concentration  (mg/L) | Concentration  (mM) | Concentration (mg/L) | Concentration  (mM) |
| **Amino acids** | | | | |
| Glycine | 30 | 0,4 | 30 | 0,4 |
| L-Arginine HCl | 84 | 0,398104 | 84 | 0,398104 |
| L-Cystine 2 HCl | 63 | 0,201278 | 63 | 0,201278 |
| L-Glutamine | 584 | 4 | 584 | 4 |
| L-Histidine hydrochloride-H_2_O | 42 | 0,2 | 42 | 0,2 |
| L-Isoleucine | 105 | 0,801527 | 105 | 0,801527 |
| L-Leucine | 105 | 0,801527 | 105 | 0,801527 |
| L-Lysine HCl | 146 | 0,797814 | 146 | 0,797814 |
| L-Methionine | 30 | 0,201342 | - | - |
| L-Phenylalanine | 66 | 0,4 | 66 | 0,4 |
| L-Serine | 42 | 0,4 | 42 | 0,4 |
| L-Threonine | - | - | 95 | 0,798319 |
| L-Tryptophan | 16 | 0,078431 | 16 | 0,078431 |
| L-Tyrosine disodium salt dihydrate | 104 | 0,398467 | 104 | 0,398467 |
| L-Valine | 94 | 0,803419 | 94 | 0,803419 |
| **MEM vitamine mix. 1x, diluted from 100x (Fisher Scientific, Cat. No.: 11120037)** | | | | |
| Choline chloride | 1 | 0,007143 | 1 | 0,007143 |
| D-Calcium pantothenate | 1 | 0,002096 | 1 | 0,002096 |
| Folic Acid | 1 | 0,002268 | 1 | 0,002268 |
| Nicotinamide | 1 | 0,008197 | 1 | 0,008197 |
| Pyridoxal HCl | 1 | 0,004854 | 1 | 0,004854 |
| Riboflavin | 0,1 | 0,000266 | 0,1 | 0,000266 |
| Thiamine HCl | 1 | 0,002967 | 1 | 0,002967 |
| i-Inositol | 2 | 0,011111 | 2 | 0,011111 |
| **Earl's balanced salt solution (EBSS, Fisher Scientific, Cat. No.: 11540616)** | | | | |
| Calcium Chloride (CaCl_2_) (anhydr.) | 200 | 1,801802 | 200 | 1,801802 |
| Magnesium Sulfate (MgSO_4_-7H_2_O) | 200 | 0,813008 | 200 | 0,813008 |
| Potassium Chloride (KCl) | 400 | 5,333333 | 400 | 5,333333 |
| Sodium Bicarbonate (NaHCO3) | 2200 | 26,19048 | 2200 | 26,19048 |
| Sodium Chloride (NaCl) | 6808,5 | 117,3879 | 6808,5 | 117,3879 |
| Sodium Phosphate monobasic (NaH_2_PO_4_-H_2_O) | 140 | 1,014493 | 140 | 1,014493 |
| D-Glucose (Dextrose) | 4500 | 25 | 4500 | 25 |
| Phenol Red | 10 | 0,025126 | 10 | 0,025126 |

**Table S2. Formulations of threonine-free medium and methionine-free medium based on Dulbecco’s modified Eagle’s medium (DMEM)**. Earl’s balanced salt solution was used as base for the custom media. Amino acids, D-glucose and MEM vitamin mix (100x) were added to the final concentrations outlined in the table. Threonine was omitted in threonine-free medium (green) and Methionine was omitted in methionine-free medium (blue)
